# Detecting microbial engraftment after FMT: sorting signals from noise using placebo sequencing and culture-enriched metagenomics

**DOI:** 10.1101/2024.09.11.612315

**Authors:** Shahrokh Shekarriz, Jake C. Szamosi, Fiona J. Whelan, Jennifer T. Lau, Josie Libertucci, Laura Rossi, Michelle Shah, Melanie Wolfe, Christine H. Lee, Paul Moayyedi, Michael G. Surette

## Abstract

Fecal microbiota transplantation (FMT) has shown efficacy for the treatment of ulcerative colitis but with variable response between patients and trials. The mechanisms underlying FMT’s therapeutic effects remains poorly understood but is generally assumed to involve engraftment of donor microbiota into the recipient’s microbiome. Previous studies have reported microbial engraftment following FMT in various disease contexts but with inconsistent results between studies. Here we investigate engraftment in UC patients receiving FMT from a single donor applying amplicon-based profiling, shotgun metagenomics and culture-enriched metagenomics. Placebo samples were included to estimate engraftment noise, and a significant level of false-positive engraftment was observed which confounds the prediction of true engraftment. We show that analyzing engraftment across multiple patients from a single donor enhances the accuracy of detection. We identified a unique set of genes engrafted in responders to FMT which supports strain displacement as the primary mechanism of engraftment in our cohort.

---

Ulcerative colitis (UC) is a type of inflammatory bowel disease (IBD) of unknown etiology that is restricted to the colon^1^. UC is believed to arise from a disruption in the balance between the immune system and microbiota in a genetically susceptible individual^2^. Current standard medical treatments have focused on suppressing the immune response and are not always effective at controlling disease^4^. An alternative approach is to alter the microbial environment responsible for driving the immune response^5^. Fecal microbiota transplantation (FMT) has emerged as an increasingly popular approach to alter the colonic microbiota^6^ and is a standard therapy for patients with recurrent *Clostridioides difficile* infection (rCDI)^7,8^ and has also been evaluated in UC^9–12^.

We reported the first randomized controlled trial (RCT) of FMT for patients with active UC^5^. This RCT showed that the proportion of patients with active UC in which remission was induced after FMT (24%) was significantly higher than a placebo group (5%), with no difference in adverse events. This has been replicated by other researchers and a systematic review of 12 RCTs suggests there is moderate quality evidence that FMT can induce remission in active UC^13^. One of the donors involved in our trial was more successful at inducing remission than other donors, with 7 of the 9 responders receiving Donor B stool^5^. Our RCT did use single donors for each participant with only a small pool of donors overall^5^. We were therefore positioned to study donor effects, and we are building on our previous work by further investigating the microbial composition of patients who received FMT from Donor B compared to those who received placebo treatments to ask whether a specific group of microbes were engrafted following FMT, and to determine whether microbial engraftment is associated with remission after FMT.

Previous studies have reported microbial engraftment—the transfer of microbes from donor to patients—following FMT in various disease contexts, particularly in rCDI^14–19^. However, there are inconsistent results between studies. Even larger studies, which applied their bioinformatic workflows to multiple publicly available datasets and disease types^20–22^, reported disparate outcomes. To our knowledge, none of these studies have applied their analytical pipelines to placebo samples from RCTs sequenced with a metagenomic approach to assess background noise or establish thresholds for engraftment detection. However, placebo samples have been used to examine background noise in a defined consortium to treat rCDI^23,24^, and alternatively, samples from the same and different individuals have been used to estimate potential noise in colonization^21,25^. Moreover, culture-independent approaches often lack the sensitivity to detect low-abundance bacteria. Culture-enriched sequencing methods provide a more comprehensive view of the human microbiome than culture-independent sequencing, particularly for low-abundance bacteria, and past studies have shown the utility of this approach in capturing the diversity of intestinal^26,27^, lung^28,29^, skin^30^, and urine^31^ human microbiota.

To answer the question of whether specific groups of microbes are responsible for inducing remission in UC, we have therefore used three high-throughput sequencing approaches; culture-independent (direct) 16S rRNA gene amplicon sequencing, culture-independent (direct) shotgun metagenomic sequencing (DMG), and culture-enriched shotgun metagenomic sequencing (CEMG). Further, we asked whether observed engraftment truly reflected the donor microbiome’s influence, or if it was an artefact of low sampling resolution. To address this, we used a unique dataset, containing stool samples from patients receiving FMT from a single donor and those procured from patients undergoing placebo treatments. This multifaceted approach enables us to provide insights into the microbiome’s potential role in UC remission.

## Results

Our RCT involved 70 patients, with 36 undergoing FMT and 34 receiving a placebo^5^. At the trial’s conclusion, 9 patients in the FMT group (7 from Donor B) and 2 in the placebo group had entered remission **(Figure 1A)**. For an accurate assessment of true microbial engraftment from Donor B, we revisited the 16S rRNA gene amplicon sequencing data with enhanced resolution by using amplicon sequence variants (ASV)^32^ instead of operational taxonomic units (OTUs). This reanalysis involved paired pre- and post-treatment samples from 20 FMT recipients of Donor B’s stool, and 31 placebo recipients **(Figure 1B).** We selected all available responders and an equal number non-responders from the Donor B FMT group (6 per group) and both responders and 10 non-responders (chosen at random) from the placebo groups for shotgun metagenomics. Additionally, shotgun metagenomic sequencing was applied to four longitudinal samples from Donor B **(Figure 1C)**. We built a comprehensive sequence library of Donor B’s samples via culture-enriched metagenomic sequencing, assembly, and annotations previously described^29^ **(Figure 1D**, see Supplementary Results**, Supp Figure 2).**

**Figure 1.**
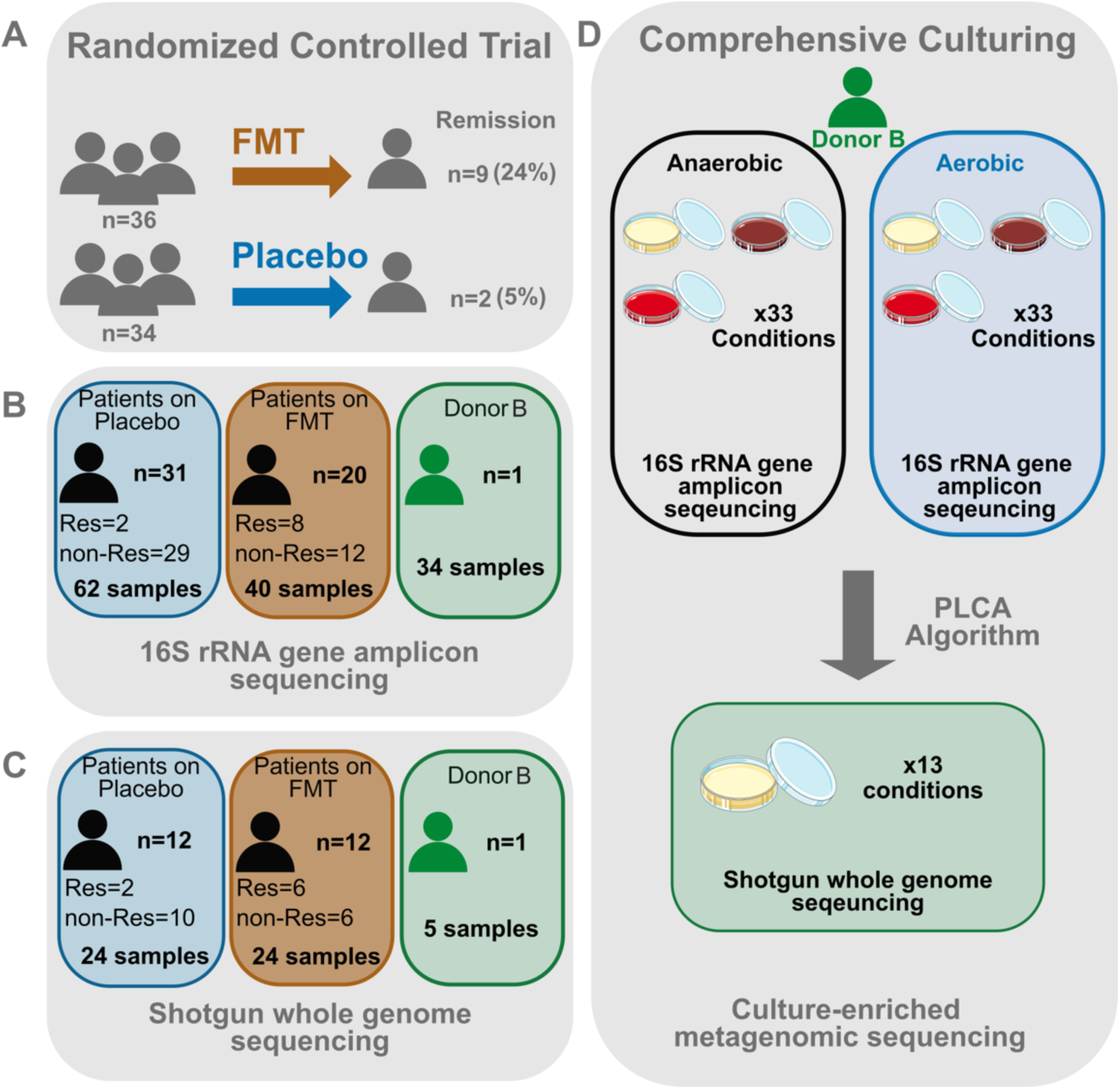
Diagram of study design. **A.** The double-blind randomized control trial of FMT for UC patients ^5^. **B.** Direct 16S rRNA gene amplicon sequencing of stool was conducted for all patients and Donor B samples. Note that no baseline sample was available for one of the Donor B responders and this subject was excluded from our analysis. **C** Shotgun metagenomic sequencing was carried out for a subset of patients who received FMT from Donor B. 6 responders and 6 non-responders and an equivalent number of placebo treated patients. **D.** The culture-enriched metagenomics workflow to build a comprehensive sequence database of Donor B’s gut microbiome. FMT = fecal microbiome transplant; PLCA = plate coverage algorithm.

**Figure 2.**
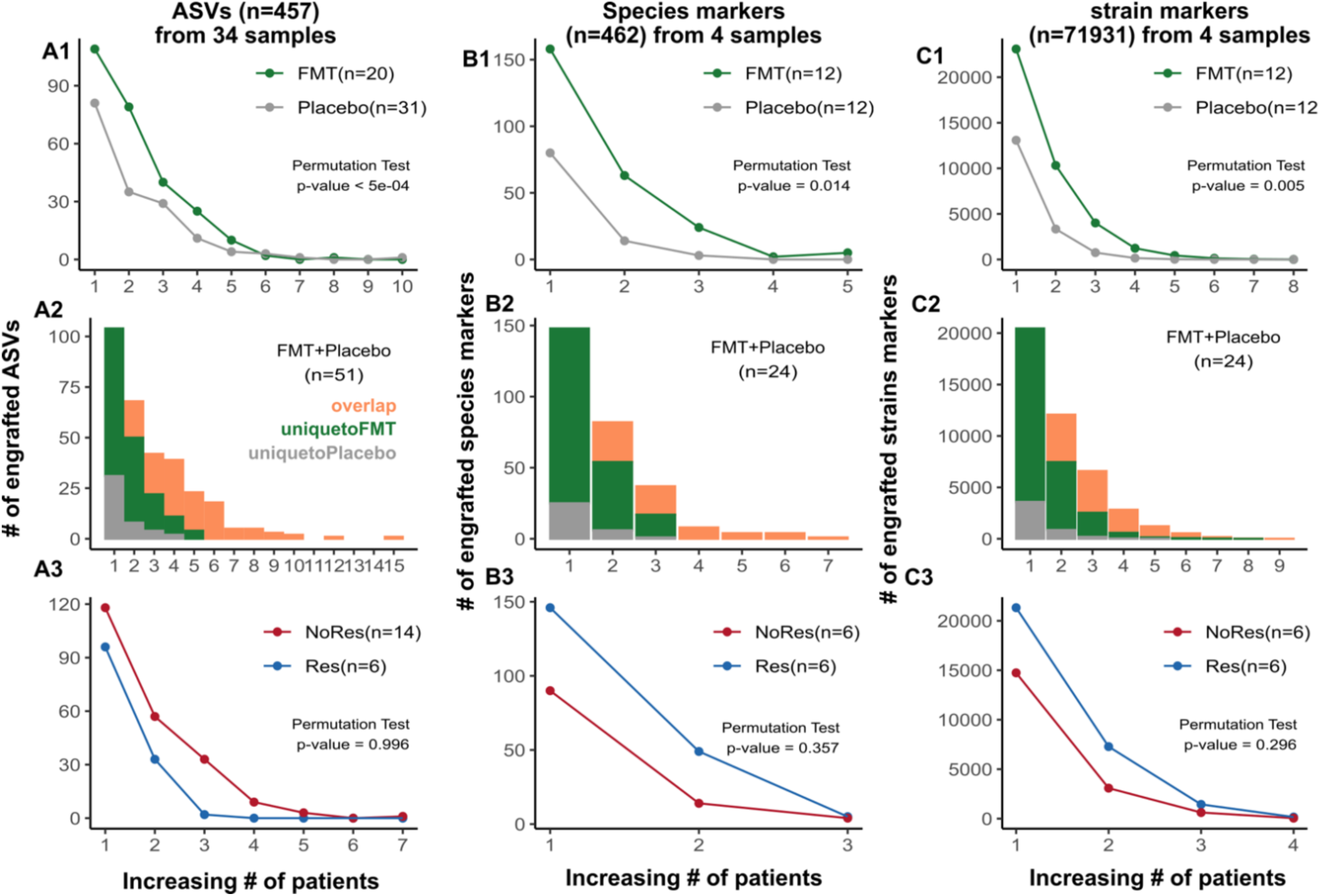
Comparison of rates of apparent engraftment of donor feature between FMT and placebo groups. Detection of apparently engrafted ASVs from 16S rRNA gene sequencing (A), species predicted from shotgun metagenomic sequencing (B) and strain markers predicted from shotgun metagenomic sequencing (C). Row 1: Comparison of the number of apparently engrafted features observed in FMT vs. placebo samples. The x-axis shows the number of patients a feature is apparently engrafted present in, and the y-axis shows the number of features observed to be apparently engrafted in exactly that many patients within FMT (green) or placebo (grey) samples. Row 2: Counting the number of apparently engrafted features that are unique to FMT samples (green), placebo samples (grey), or present in both (orange). The x-axis shows the number of patients a feature is apparently engrafted in, and the y-axis shows the number of features observed apparently engrafted in exactly that many patients. Row 3: The number of apparently engrafted features in FMT samples that are present in patients who responded to FMT treatment (blue lines) and did not respond (red lines). The differences between the points on the green and grey (row 1) or blue and red (row 3) lines were used to calculate the test statistics for the permutation tests.

### Microbial engraftment is present after FMT against a background of false positives

We have previously shown that culture-independent methods often underestimate microbial diversity in a stool sample compared to culture-enriched approaches^26^. If a patient has a donor feature below the detection limit at baseline but it is detected after FMT, this would appear as engraftment but would be a false positive. We refer to this as spurious engraftment. Here we compare measurements of engraftment in FMT recipients from a single donor and placebo recipients to estimate the extent of spurious engraftment at different sequencing resolutions, including 16S rRNA gene amplicons (ASVs) , short-read metagenomics analysis at species (MetaPhlAn4^33^), and strain ( StrainPhlAn4^33^) levels.

Conceptually, there should be no engraftment in the placebo group, and we hypothesized that spurious engraftment (false positives) would be minimal. Furthermore, we expect significant microbial engraftment to correlate with clinical remission after FMT. To investigate this, we utilized samples from Donor B in both 16S rRNA and shotgun metagenomics sequencing datasets, establishing a stringent cut-off to define apparent microbial engraftment. A microbe was only described as engrafted if its relative abundance was zero prior to FMT and rose above our established cut-off after FMT (detailed in Methods, **Supp Figure 1**).

Initially, we investigated whether a clear difference exists between rates of apparent engraftment in the FMT and placebo groups. Our dataset, involving a single donor, allowed us to identify features that were apparently engrafted in multiple patients. We postulate that engraftments observed in multiple individuals are more likely to be the result of true microbial colonization and less reflective of individual-specific microbial changes during FMT. We identified donor ASVs, species, and strain markers that appeared to be engrafted in a given numbers of patients ranging from one, two, three, etc. from both FMT and placebo groups **(Figure 2 A1, B1, and C1)**. We then computed the differences in numbers of each type of engraftment event for each dataset (donor vs placebo) and used permutation tests to determine whether the amount of apparent engraftment observed in patients who received FMT was significantly different from the amount observed in placebo (see Methods). The results showed that the FMT group had a significantly higher number of apparent engraftments than the placebo group (p-values presented in **Figure 2** for each dataset), highlighting the likelihood that FMT leads to (at least some) true engraftment. However, a high number of spurious engraftments were also observed in the placebo group at ASV, species, and strain marker, even with our stringent cut-off, suggesting considerable noise (false positive) in microbial engraftment detection.

Next, we identified apparently engrafted donor ASVs, species, and strain markers that were unique to either the FMT or placebo groups, or shared between them (“overlap”), as shown in **Figure 2 A2**, **B2**, and **C2**. The shotgun metagenomic dataset contained a stronger signal than the 16S dataset, showing that a higher proportion of apparently engrafted features are unique to the FMT group. Interestingly, the number of apparent engraftments in the unique-to-placebo group that were shared in more than one individual was reduced in species- and strain-level analysis. This indicates that looking for features that appear to engraft in multiple patients reduces the risk of detecting spurious engraftment. The number of apparently engrafted features that were observed in both FMT and placebo groups underscores the importance of including placebo sequencing in studies, and reflects the fact that common microbial changes in individuals over time can mimic an engraftment signal.

Last, we explored whether microbial engraftment is associated with the response to FMT treatment. We compared the number of apparently engrafted features in responder and non-responder patients in the FMT group across ASV, species, and strain marker datasets **(Figure 2 A3, B3, C3)**. The 16S dataset included 14 non-responders and 6 responders, while the shotgun metagenomic datasets included the same 6 responders and a sub-set of non-responders (n=6). Similar to the placebo vs. FMT comparisons described above, we calculated the differences in the number of engrafted features and weighted engraftment statistics (see Methods) and used permutation tests to compare the two groups. The results indicated no clear difference in the number of apparent engraftments between responders and non-responders. Numbers of apparently engrafted features can be found in **Supplementary Table 2** and **3**.

### High resolution mapping of patient reads to Donor B MAGs to detect apparent genome engraftment

The results above indicate that detection of microbial engraftment may require greater microbial resolution than amplicon-based or read-based shotgun metagenomic sequencing. We hypothesized that co-assembling sequences from direct and culture-enriched metagenomic sequencing (CEMG) of the Donor B microbiota would improve gene and metagenome assembled genome (MAG) recovery from this donor compared to the commonly used direct metagenomic (DMG) approach. To generate the highest possible resolution assembly of Donor B’s microbiota, we built a comprehensive database using a co-assembly of four longitudinal DMG samples as well as our single CEMG sample (see Supplementary Results, **Supp Figure 2**). We focused on 209 high quality MAGs out of a total of 447 bins in our databases. To track the engraftment of these MAGs after FMT, we mapped raw shotgun reads — subsampled to uniform sequencing depth — from the pre- and post-treatment samples of 24 patients (12 FMT recipients and 12 placebo-treated individuals, 48 samples in total) to these 209 MAGs.

To define the thresholds for MAG engraftment, we mapped raw sequence reads from our four Donor B samples to the Donor B MAG database. In each sample, we observed bimodal distribution in MAG coverage, with the modes at the extremities of the distribution. We reasoned that MAGs that were present in that sample were the ones that appeared in the upper mode, where at least 75% of the length of the MAG was covered at least 1x, and the MAGS that were absent from that sample were those that appeared in the lower mode, where no more than 20% of the length of the MAG was covered at least 1x. We therefore set these coverage values as our cutoffs for presence and absence, respectively (**Supp Figure 1**).

We assessed the number of apparently engrafted MAGs in FMT recipients compared to placebo recipients, using methods similar to those in the read-based metagenomic analyses described above. Our goal was to determine if assembly-based assessments would reveal clear differences between the FMT and placebo groups. Consistent with previous results, we found a significant increase in the number of MAGs apparently engrafted in any given number of patients in the FMT group compared to placebo (**Figure 3A1**, permutation test p=0.008). We observed no significant differences in the numbers of engrafted MAGS, or the numbers of patients per engrafted MAG, when comparing FMT responders to non-responders (**Figure 3A3**, permutation test p=0.24).

**Figure 3.**
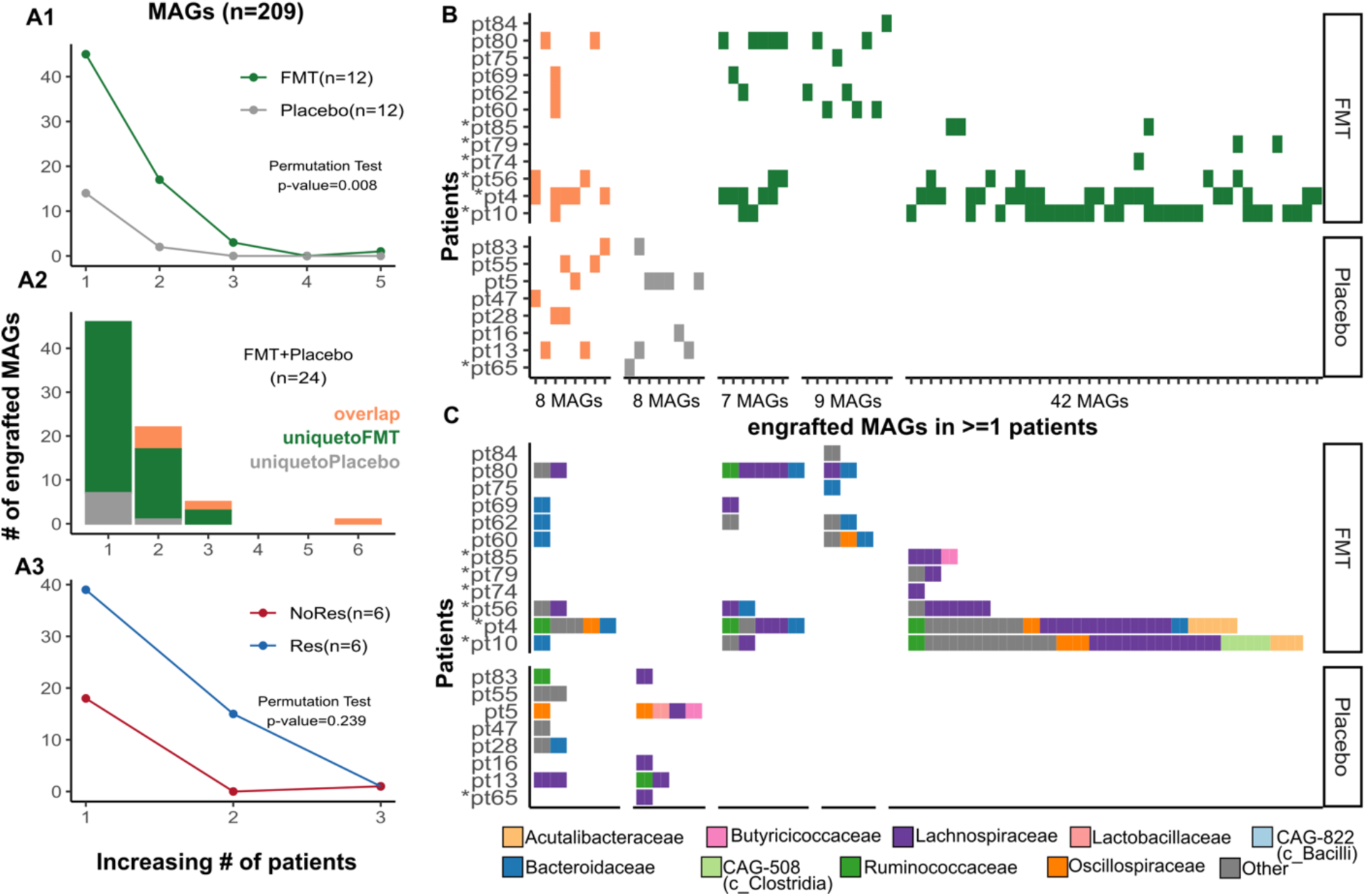
High resolution genome-resolved metagenomics reveals apparent microbial genome engraftment following FMT. **A1.** Comparison of apparently engrafted MAGs in patients who received FMT (n=12) vs. placebo (n=12) treatment. The x-axis displays number of patients in which a given MAG appears engrafted. **A2.** Counts of MAGs engrafted in only FMT (green), only Placebo (grey), and both (orange). **A3.** MAGs engrafted in FMT patients, observed in responders (n=6, blue) and non-responders (n=6, red). The differences between the points on the green and grey (A1) or blue and red (A3) lines were used to calculate the test statistics for the permutation tests. **B.** All donor MAGs engrafted in at least one patient. Green represents MAGs exclusively engrafted in FMT recipients, grey indicates MAGs spuriously detected as engrafted in the Placebo group, and orange signifies MAGs detected in both FMT and Placebo treatment cohorts. MAGs are ordered on the x axis by which group(s) of patients they were apparently engrafted in. **C.** The family-taxonomy of donor MAGs engrafted in at least one patient.

Upon examining MAGs engrafted in one or more individuals, we found that 57% of engrafted MAGs (42 out of 74) were unique to patients who received and responded to FMT (**Figure 3B**), and most of these events were observed in 3 recipients (y patients 4, 10, and 56). *Lachnospiraceae*, *Ruminococcaceae*, and *Oscillospiraceae* were the most abundant families among these MAGs (**Figure 3C**). These findings suggest that microbial genomic alterations in response to FMT are individual-specific, and that a degree of spurious engraftment is observable in the placebo group, indicating that even genome-resolved metagenomic approaches to detect engraftment are also prone to false positives.

### A signature of gene engraftment in patients who responded to FMT

Although culture enrichment allowed us to refine 209 MAGs from a single FMT donor, tracking MAGs omits information from the majority of assembled reads, which remain in bins or contigs excluded from MAGs. Since MAGs represent incomplete genomes^34^, a more comprehensive approach to investigating engraftment would focus on all donor genes.

We aimed to identify genes linked with engraftment and response to FMT. Using all Donor B contigs >2.5kb long, we identified a total of 755,662 genes. We then used precise mapping of raw reads from all the patient samples (n=48), to the Donor B gene database. We define apparently engrafted genes as those with ≤ 1x coverage over less than 25% of their length in the baseline sample, and ≥ 1x coverage over 90% or more of their length after treatment.

We employed the same methodology used in **Figure 2** to assess engraftment in the FMT and placebo groups. Our results revealed a significant difference in the number of apparently engrafted genes between patients treated with FMT and those receiving placebo (**Figure 4A1**, permutation test p=0.001). Consistent with all previous analyses, the number of apparently engrafted genes unique to the placebo group decreased as we restricted the count to genes apparently engrafted in increasing numbers of patients (**Figure 4A2**). Consistent with other datasets, we could not clearly see a difference in the number of engrafted genes between responders and non-responders within the FMT group (**Figure 4A3,** permutation test p=0.86).

**Figure 4.**
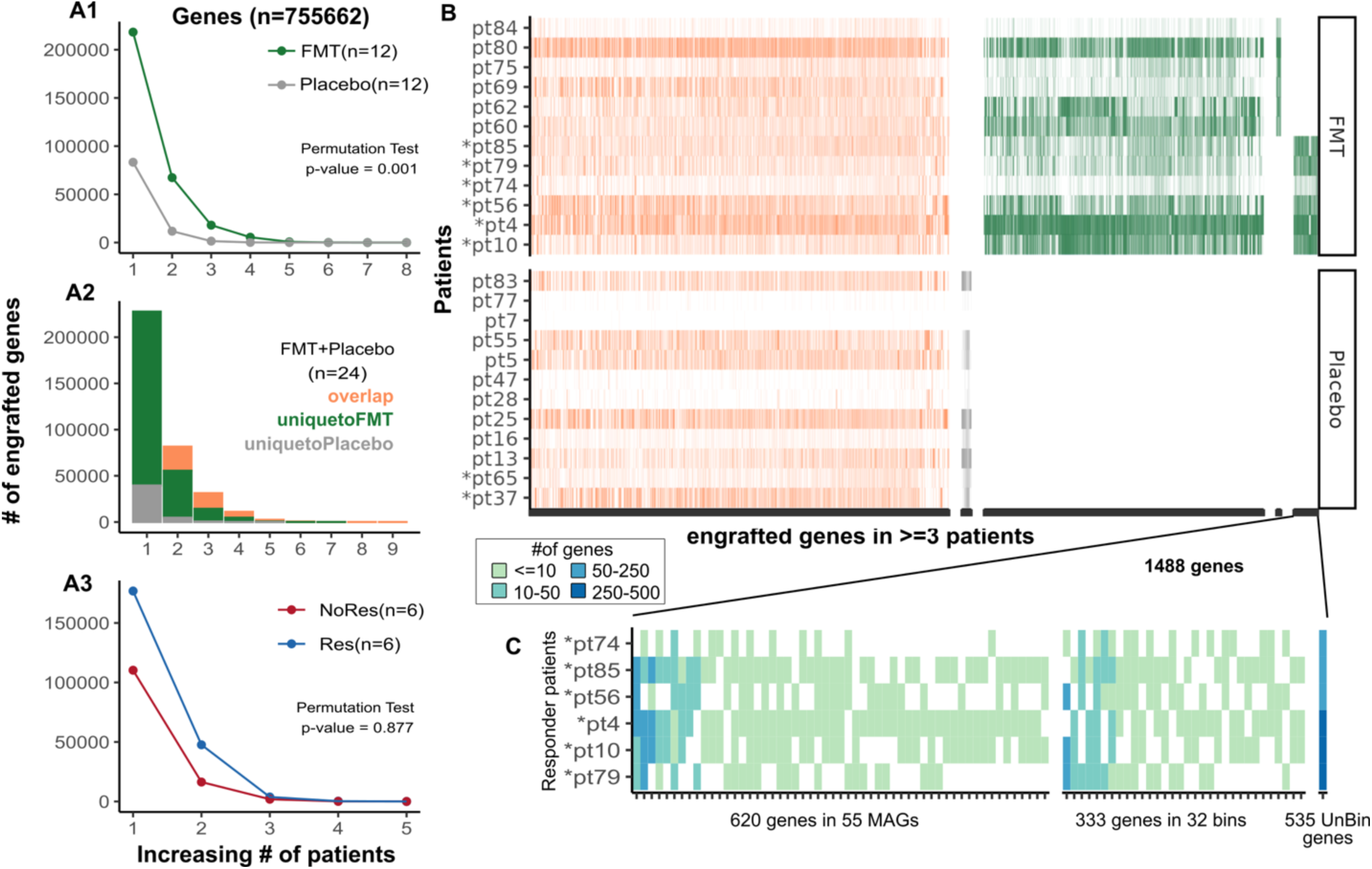
Identifying genes commonly engrafted in FMT responders. **A.** Counts of apparently engrafted genes in FMT and placebo groups. **A1.** Examination of apparently engrafted genes in patients receiving FMT (n=12) or placebo (n=12) treatment. The x-axis shows the number of patients in which each gene appears to be engrafted, emphasizing common gene engraftment. **A2.** Discriminating donor gene engraftment patterns across all patients (n=24) to distinguish unique engraftment in FMT (green), in Placebo (grey), and genes engrafted in both FMT and Placebo (orange). **A3.** Comparison of apparently engrafted genes in FMT patients in responders (n=6, blue) and non-responders (n=6, red). The differences between the points on the green and grey lines (A1) or blue and red lines (A3) were used to calculate the test statistics for the permutation tests. **B.** Donor genes that engrafted in at least 3 patients are categorized as follows: genes engrafted in FMT patients but also present in the placebo group are shown in orange; genes unique to the placebo or FMT groups are depicted in grey and green, respectively. Among genes apparently engrafted only in FMT, those found in both responders and non-responders, as well as those unique to each group, are separately grouped on the x-axis. **C.** Location of the 1,488 genes apparently uniquely engrafted in 3 or more FMT responders in our Donor B assemblies. Among the 1,488 genes, 333 were found in 32 metagenomic bins, 620 were present in 55 MAGs, and 535 genes were located on un-binned contigs (“UnBin”).

To enrich for legitimate engraftment events, we focused on genes engrafted in three or more patients. This provided a set of 45,419 genes. Among these, 56.9% (25,826) were engrafted in both FMT and placebo recipients (Figure *4*B, orange). Notably, 37.9% (17,230) were exclusive to FMT recipients (Figure *4*B, green), while only 5.2% (2,363) were unique to the placebo group (Figure *4*B, grey). This final category, since it is known to be false positives, was not subject to further analysis. Genes unique to responders represented 3.3% (1,488), compared to only 0.6% (283) in non-responders (**Figure 4B**). To trace the origins of genes uniquely associated with FMT responders, we examined their presence in contigs grouped in MAGs and non-MAG bins. We found 620 genes within 55 unique MAGs, 535 in contigs not assigned to any bin (“UnBin”), and 333 in 32 non-MAG bins (**Figure 4C**). These findings highlight the importance of our gene-based approach. Relying solely on genome detection strategies using MAGs would have prevented the identification of most of these genes.

### Taxonomic specificity of commonly engrafted genes and distribution in an independent IBD patients

To further investigate the 1,488 genes engrafted in at least 3 responders, we searched for these genes in 33,167 RefSeq genomes from all bacterial phyla, updated as of May 2023. Our objective was to ascertain whether these genes are widely distributed across various taxa, or are specific to certain strains.

Our analysis revealed that 66 out of the 33,167 genomes contained the genes of interest, each with more than five genes per genome. These genomes were distributed across six bacterial families (**Figure 5A**). Notably, genes identified in the families *Bacteroidaceae* were present in multiple species within their genera, suggesting these were widely shared between strains within the *Bacteroides* genus. By contrast, genes present in *Lachnospiraceae* and *Ruminococcaceae*, were more strain-specific within a species. The remaining families contained 11% of the detected genes. (**Figure 5B**).

**Figure 5.**
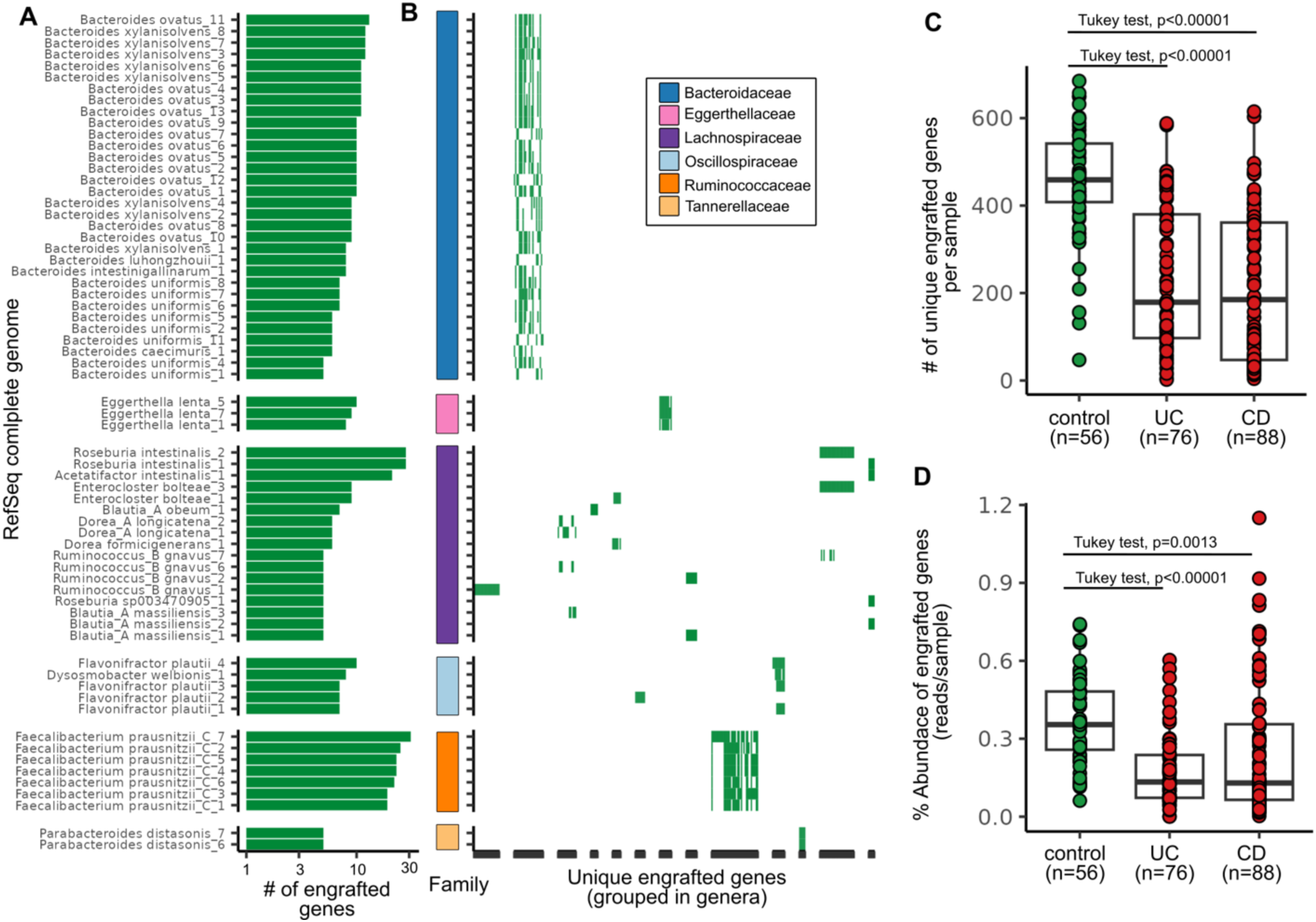
Commonly engrafted genes in FMT responders are strain-specific and depleted in IBD patients. **A.** We searched for genes commonly apparently engrafted in FMT responders in the RefSeq complete bacterial genomes (n=33,167) database. The number of observed engrafted genes per genome is represented by green bars on the x-axis, while strains with at least 5 engrafted genes are displayed on the y-axis. Bacterial strains are grouped and color-coded on the y-axis according to their family. **B.** To demonstrate the specificity of these genes within a genus, unique engrafted genes are represented on the x-axis and compared across the genomes depicted on the y-axis (shared with panel A). **C.** Comparison of the number of genes commonly engrafted in responders that were found in healthy individuals (n=56) and patients with UC (n=76) and CD (n=CD). We utilized metagenomic reads from a publicly available dataset (PRJNA40007229^35^) that includes samples from healthy donors and IBD patients to explore the association of these genes with health status. **D.** Comparison of the relative abundance of these commonly engrafted genes within the same cohort as panel C.

Among the 1,488 genes identified, 1,415 were annotated as coding sequences (CDS) with a median length of 491 bp. Of these, 845 proteins were previously characterized or had predicted function by homology, while the remaining were hypothetical proteins with no predicted function. Among the characterized proteins, 593 were distributed across 19 diverse Clusters of Orthologous Groups (COG) categories. The most prevalent categories included Transcription, which accounted for 12.8% of these proteins, followed by Replication, Recombination, and Repair (12.3%), and Carbohydrate Transport and Metabolism (8%).

One notable observation (**Supp Figure 3**) is that for any individual only a small portion of the microbiome features are shared between patient and donor. However, most of the donor’s microbiome features are shared with at least one subject. This is also true of engrafted features. This reflects the high individual variability in human gut microbiomes.

Expanding our investigation, we utilized a larger dataset of publicly available metagenomic samples (PRJNA40007229^35^), which included healthy individuals (n=56), UC patients (n=76), and patients with Crohn’s disease (CD) (n=88). We mapped these reads to the 1,488 genes to determine their presence and relative abundance in the samples. Intriguingly, we found a notable reduction in both the number and relative abundance of these genes in IBD patients compared to healthy individuals. (Tukey test, p-values in **Figure 5C and 5D**, overall ANOVA p-values <0.0001). These observations, derived from an independent and more extensive dataset, further underscore the potential significance of these genes in the context of IBD.

## Discussion

FMT has emerged as a promising therapeutic option for UC, demonstrating efficacy in RCTs when compared to placebo^13,36^. However, the variability in patient response remains a critical challenge, underscoring the need to understand the factors influencing the success of FMT. Donor microbiota composition has been implicated in treatment outcomes, with studies reporting both donor-dependent efficacy and improved outcomes when pooling microbiomes from multiple donors^5,18,37^. In this study, we aimed to dissect the mechanisms underlying successful FMT by examining microbial engraftment in UC patients.

We examined the rates of apparent engraftment in both FMT and placebo groups. We observe a high rate of spurious engraftment (false-positives) in the placebo group which confounds the prediction of true engraftment. This demonstrates that it is not sufficient for a feature to meet engraftment criteria of absence in a baseline and presence in both donor and post-FMT samples. Additionally, our data suggest that analyzing engraftment across multiple patients from a single donor enhances the accuracy of detection. It is possible that placebo engraftment represents the acquisition of strains from environmental sources. However, the probability that environmental sources of spurious engrafted features match donor is low.

Our analysis also underscores the necessity of integrating placebo group sequencing in investigation of post-FMT engraftment. Our data from the placebo controls demonstrated that a substantial amount of false positives detection occurs, and caution must be taken when interpreting results. These false-positives arise because donor features in a patient (e.g. species/strains/MAGs/genes) may be present at baseline but below the level of detection. If these features increase and are detected at a second timepoint, then they will appear to be engrafted. By refining engraftment thresholds and employing rigorous controls, we aim to provide a clearer and more reliable framework for interpreting FMT outcomes in UC. This is not unexpected since even deep sequencing does not capture the full complement of microbes present. As we have shown previously^26,28,29^ and here **(Supp Figure 2)**, comprehensive culture methods enrich the number of features detected 3-4 fold.

The application of MAGs in studying microbial engraftment after FMT presents significant challenges. The inherent difficulties in accurately assembling MAGs from complex microbial communities result in the exclusion of a substantial portion of sequencing data (>50%). The non-MAG metagenomic data will include species present at lower abundance, plasmids, mobile genetic elements, and when multiple strains of a species are present - strain-specific genes. Using the define thresholds for donor MAG presence and absence in subjects, we observed 4 MAGs predicted to be engrafted in at least three FMT recipients, but only one of these, a *Lachnospiraceae*, was shared between 3 responders (*Hominisplanchenecus faecis*).

The clearest signals of engraftment in our study came from the gene dataset where specific sets of donor genes were predicted to be engrafted and associated with response in multiple FMT recipients. The gene-level analysis is consistent with most engraftment events being associated with strain displacement where a donor strain replaces a recipient strain of the same species. It is also possible that some engrafted genes may represent horizontal gene transfer events. Here we use strain in the context of the presence and absence of accessory genes which focuses on functional differences between strains. Approximately 60% of these genes were not found in MAGs even though our culture-enriched metagenomic increased the number of MAGS 4-fold and increased the size of most MAGs compared to standard metagenomics. Furthermore, MAGs containing these genes (n=55) were not predicted to be engrafted. This reflects the incomplete nature of MAGs and the challenges of establishing the thresholds set for presence and absence of features (**Supp Figure 1**).

Strains can be defined by single nucleotide polymorphisms in defined set of single core genes (e.g. StrainPhlAn). This is a powerful approach for tracking lineages over time and other evolutionary studies as it avoids the confounding effects of horizontal gene transfer. The strain-sharing inference pipeline within StrainPhlAn calculates pairwise distances between samples without distinguishing unique strain properties. Applying this method to our data, we found a strong signature of engraftment (**Suppl Fig 7**) and only a small number of the total species-level bins passed the threshold to be included 54 out of 252, **Supp Figure 8**. However, this is a distance matrix of the distribution of strain markers within a species and it does not identify specific engrafted strains per se. So while the results suggest that engraftment is occurring within a species, we can not identify a specific strain or marker associated with this. Other limitations of the strain-sharing inference pipeline for our specific question of identifying engrafted features are discussed in the supplementary methods.

Using a gene-centric methodology we identified specific sets of genes uniquely present in patients who responded to FMT treatment. Many of these genes are involved in diverse microbial metabolic pathways. However, a main challenge in interpreting these results arises from the limited annotations available in existing databases, and a significant proportion of the predicted genes (∼25%) have no predicted function. Our study was limited by small number of subjects and responders in our trial. Despite these limitations, when examining these commonly engrafted genes with a larger cohort of IBD patients, we found that they were consistently depleted, suggesting a possible protective or beneficial role for these genes that is disrupted in this disease state.

Our analysis revealed a notable pattern of strain specificity among these genes, particularly in species within the *Lachnospiraceae* and *Ruminococcaceae* families. Interestingly, the same taxa were among the top engrafted bacteria in other feature types (**Supp Figure 9**). This specificity suggests that certain strains of these species may be important for achieving the desired therapeutic outcomes of FMT. Conversely, in the *Bacteroides* genus, we observed that these genes are not confined to strains of specific species but are rather widespread across the genus (**Figure 5**). As might be predicted from this distribution, there is relatively low level of prediction for *Bacteroides* MAG engraftment (**Figure 3**). This distribution pattern could indicate a broader functional role for these genes within the *Bacteroides*, or may be reflective of an accessory genome that is more widely shared across the family.

Strain level engraftment may also reflect direct strain-strain competition where donor strains that are more adept at outcompeting a patient’s own strains and may be independent of response to FMT. However, we do find donor genes specifically engrafted in multiple responders, and also show these genes are depleted in stool metagenomic samples from ulcerative colitis and Crohn’s disease patients compared to healthy controls (**Figure 5C,D**). These observations provide support for a role of these genes in response to FMT.

The objective of this study was to attempt to identify engrafted features of the donor that were associated with clinical response in patients. The high level of false positive engraftment in the placebo group highlights one of the challenges in studying engraftment. This was independent of the methodological approach applied and is an inherent challenge amplicon of metagenomics studies of microbiomes. An interest in this field has been to replace FMT with defined consortia of microorganisms. For reasons discussed above, defining strains based on precise gene level engraftment may more accurately reflect functional requirements for response. A consortium of strains defined by gene-level engraftment might be more effective as a FMT alternative than a consortia based only on species considerations. By focusing on these key microbial players and their functional genes, and investigating their roles in health and disease, we can begin to piece together the intricate mechanisms through which the microbiota exerts their beneficial effects, paving the way for refined FMT strategies tailored to individual microbial compositions.

## Methods

### Study design and sample collection

The study design and sample collection was as described previously^5^. Briefly, 70 active UC patients (Mayo score >=4 with an endoscopic Mayo score >=1) were randomly assigned to either 6 weeks of FMT (once per week; 50 mL, via enema, from healthy anonymous donor) or placebo (once per week; 50 mL water enema) in a double-blind randomized controlled trial. Stool samples were collected at baseline (before treatment), and during each week of the trial.

### DNA extraction and 16S rRNA gene sequencing

Genomic DNA extraction and PCR amplification of the V3 region of 16S rRNA gene was conducted using previously described protocols^5,38,39^. Briefly, 0.2 g of fecal matter was mechanically homogenized using ceramic beads in 800 μL of 200 mM NaPO 4 (pH 8) and 100 μL of guanidine thiocyanate-EDTA-N-lauroyl sacosine. This was followed by enzymatic lysis of the supernatant using 50 μL of 100 mg/mL lysozyme, 50 μL of 10 U/μL mutanolysin, and 10 μL of 10 mg/mL RNase A for one hour at 37 °C. Then, 25 μL of 25% sodium dodecyl sulfate (SDS), 25μL of 20 mg/mL proteinase K, and 75 μL of 5 M NaCl was added, and incubated for one hour at 65 °C. Supernatants were collected and purified through the addition of phenolchloroform-isoamyl alcohol (25:24:1; Sigma, St. Louis, MO, USA). DNA was recovered using the DNA Clean && Concentrator TM −25 columns, as per manufacturer’s instructions (Zymo, Irvine, CA, USA) and quantified using the NanoDrop (Thermofisher, Burlington, ON). After genomic DNA extraction, the V3 region of the 16S rRNA gene was amplified via PCR using these conditions per reaction well: Total polymerase chain reaction volume of 50 μL (5 μL of 10X buffer, 1.5 μL of 50mM MgCl 2 , 1 μL of 10 mM dNTPs, 2 μL of 10mg/mL BSA, 5 μL of 1 μM of each primer, 0.25 μL of Taq polymerase (1.25U/ μL), and 30.25 μL of dH 2 O. Each reaction was divided into triplicate for greater efficiency. The primers used in this study were developed by Bartram et al.,2011. PCR conditions used included an initial denaturation at 94 °C for 2 minutes, followed by 30 cycles of 94 °C for 30s, 50 °C for 30s, 72 °C for 30s, followed by a final elongation at 72 °C for 10 minutes. All samples were sequenced using an Illumina MiSeq platform at the McMaster Genomics Facility (Hamilton, Ontario, Canada). Samples were processed in batches, meaning not all samples were extracted and sequenced at the same time.

### 16S rRNA gene sequencing processing pipeline

Cutadapt^40^ v1.14 was used to filter and trim adapter sequences and PCR primers from the raw reads, using a quality score cut-off of 30 and a minimum read length of 100 bp. We used DADA2^32^ v1.14.0 to resolve the sequence variants from the trimmed raw reads as follow. DNA sequences were trimmed and filtered based on the quality of the reads for each Illumina run separately. The Illumina sequencing error rates were detected, and sequences were denoised to produce ASV count table. The sequence variant tables from the different Illumina runs were merged to produce a single ASV table. Chimeras were removed and taxonomy was assigned using the DADA2 implementation of the RDP classifier against the SILVA database^41^ v1.3.2, at 50% bootstrap confidence.

The ASV, taxonomy, and clinical tables were all merged into one data object in R v4.2.0 using Phyloseq v1.40.0 package^42^.

### Library preparation for shotgun metagenomic sequencing

We conducted direct shotgun metagenomic sequencing on 48 samples collected from 24 patients (12 patients who received FMT (6 responders and 6 non-responders) and 12 who received placebo (both responders and 10 non-responders)), at 2 time points each (baseline and 6 weeks after treatment), as well as 4 samples from Donor B. Genomic DNA was standardized to 5 ng/μL and sonicated to 500 bp. Using the NEBNext Multiplex Oligos for Illumina kit (New England Biolabs), DNA ends were blunted, adapter ligated, PCR amplified, and cleaned as per manufacturer’s instructions. Library preparations were sent to the McMaster Genome Facility and sequenced using the Illumina HiSeq platform, with a mean depth of approximately 18 million paired-end reads per sample.

### Culture-enriched and independent metagenomics on Donor B samples

A fresh, anaerobic fecal sample was collected from Donor B. The collected sample was cultured using 33 different media, and incubation of plates anaerobically and aerobically resulted in 66 culture conditions for culture-enriched molecular profiling using a previously described protocol^26^. The list of media and culture conditions are described therein. 16S rRNA gene amplicon sequencing was conducted on plate pools of all 66 culture conditions. To determine a subset of plates that adequately represent the sample, the distribution of ASVs in the direct sequencing was compared to the culture-enriched sequencing using the PLCA algorithm^29^. Shotgun metagenomics was conducted on the 13 plate pools identified by the PLCA algorithm as representing the community. Genomic DNA was isolated from the all 13 selected plate pools and shotgun metagenomics conducted as previously described, with a mean depth of approximately 14 million paired-end reads per plate pool. Direct shotgun metagenomics was also conducted on the same fecal sample directly.

### Comparison of the culture-enriched metagenomics with direct metagenomics data

To build a culture-enriched metagenomic library, raw shotgun sequences from the selected plate pools and the original fecal sample collected from Donor B for culturing were co-assembled as follows: Low-quality reads and sequencing primers were removed using Trimmomatic^43^ v0.38 with ‘LEADING:3 TRAILING:3 SLIDINGWINDOW:4:15 MINLEN:36’ option. The reads were decontaminated for any human DNA using DeconSeq^44^ v0.4.3. These cleaned reads were then co-assembled using metaSPAde^45^ v3.10.1. Filtered raw reads were mapped to contigs (minimum length of 2.5 kb) using BWA-mem^46^ v0.7.17, and then binned with Metabat2^47^ v2021. This was done for both the CEMG assembly and direct metagenomic sequencing (DMG) reads from the fecal sample, using identical methods. These datasets are referred to as CEMG and DMG, respectively.

The microbial compositions of DMG and CEMG datasets were then comprehensively evaluated using the following procedure: The single-copy core genes were identified within each bin using CheckM^48^ v1.1.2, any bin with minimum 70% completion and maximum 10% contamination was defined as a metagenome assembled genome (MAG). The shotgun reads were mapped to the assembled contigs to estimate sequence coverage for all contigs, Bins, and MAGs. We used BWA-mem^46^ to map reads to assembled contigs and the anvio pipeline^49^ to normalize the coverage to depth of sequencing. Detection values were calculated for each bin using anvio^49^. A detection value is defined as the proportion of a given MAG that is covered by at least one read; in other words, it estimates the proportion of MAG that recruited reads to it.

For taxonomic assignment of the metagenome-assembled genomes (MAGs), we utilized GTDB-Tk^50^ v202.0. We built a maximum-likelihood phylogenetic tree using the aligned protein sequences from GTDB with FastTree^51^. To compare the lengths of MAGs in DMG and CEMG, we selected MAGs based on their proximity in the phylogenetic tree. The total assembly length was determined by summing contig lengths, calculated with metaSPADE, in R. All figures were created and visualized using R version 4.0.3.

### Shotgun metagenomic sequencing processing pipeline

Raw reads were filtered to remove low-quality sequences and decontaminate human-derived reads using KneadData. These filtered reads were then analyzed to identify species- and strain-level markers with MetaPhlAn4 and StrainPhlAn4, respectively. To profile the relative abundance of species, MetaPhlAn4^33^ v4.0.4 was employed with the ’-t rel_ab’ option. MetaPhlAn4 introduces the ability to perform strain-level profiling using non-aggregate marker information from the StrainPhlAn4 pipeline. This method enables strain tracking and comparison across multiple samples. For strain tracking in this study, we used the ’-t marker_ab_table’ option alongside the ’--nreads’ parameter to specify the number of metagenomic reads. A default minimum Reads Per Kilobase (RPK) of 1 was applied to identify a marker as present.

We estimated strain-sharing inferences using a previously published workflow (Valles-Colomer et al., 2023^25^) based on StrainPhlAn4 v4.0.4^33^. Species-level markers, previously generated during the MetaPhlAn4 step, were extracted from all samples using the ’sample2markers.py’ script. These markers, referred to as Species-level Genome Bins (SGBs), were used to infer strain-level variation.

For every species (SGB) detected in Donor B samples (n = 4), a database containing the marker genes was built using the ’extract_markers.py’ script. Single Nucleotide Variant (SNV) profiling was performed for each Donor B species to generate phylogenetic trees using StrainPhlAn4. To enhance sensitivity for strain engraftment detection, we applied a set of predefined parameters (’--sample_with_n_markers 20’, ’--marker_in_n_samples 50’) to retain more samples and markers during SNV alignment. Pairwise phylogenetic distances for SGBs passing the SNV profiling step were calculated using the ’tree_pairwisedists.py’ script.

We utilized pre-computed species-specific strain identity thresholds from Valles-Colomer et al. (2023), which were based on comparisons of inter- and intra-individual phylogenetic distances. These thresholds, publicly available on the MetaPhlAn GitHub repository, were used to identify strain engraftment events.

Filtered metagenomic reads from donor samples (n=4) and culture-enriched plate pools (n=13) were co-assembled using metaSPAdes^45^ v3.10.1 to create a comprehensive Donor B database. Contigs of at least 2.5 kb were annotated with Bakta^52^ v1.5.0 to identify genes. These annotated contigs were then binned with Metabat2^47^ v2021, leveraging mapping data from all plate pools and Donor B samples. The quality of the resulting bins was evaluated using CheckM^48^ v1.1.2; bins with a completion rate of at least 70% and contamination of 10% or less were designated as metagenome-assembled genomes (MAGs). Taxonomic classification of MAGs was performed using GTDB-Tk^50^ v202.0.

All patient metagenomic samples (n=48) were normalized to the lowest sequencing depth observed in the dataset (5.5 million paired-end reads) using seqtk^53^ v1.1.0 for random subsampling. These rarefied reads were mapped to the genes and contigs assembled from Donor B using BWA-mem^46^ v0.7.17, applying perfect match filtering options ’-O 60 -E 10 -L 100’. Coverage information at 1x was calculated using samtools^54^ ’coverage’.

A database of 33,167 RefSeq bacterial genomes, categorized as ’assembly-level complete’ and downloaded as of May 2023, was compiled using a custom Python script. We identified a unique set of genes present exclusively in responder patients and used these as query sequences for BLAST searches against this database, employing BLAST^55^ v2.13.0+. Hits were selected based on stringent criteria: percent identity of at least 90%, query coverage of at least 90%, and gene lengths of 100 base pairs or more. Genomes containing at least five such genes were visualized. The taxonomy of these genomes was determined using GTDB-Tk^50^ v202.0. To functionally annotate these genes, they were clustered at 90% protein identity using MMseqs2^56^ v13.45111 and then profiled against KEGG, COG, and PFAM databases using eggNOG-mapper^57^ v 2.1.7 and database version of 5.0.2.

Additionally, a publicly available metagenomic dataset (PRJNA40007229^35^), comprising samples from 56 healthy subjects, 76 patients with ulcerative colitis (UC), and 88 patients with Crohn’s disease (CD), was downloaded from the Sequence Read Archive (SRA). These samples were mapped to the identified gene set using BWA-mem^46^ v0.7.17, applying strict matching filters (’-O 60 -E 10 -L 100’). samtools^54^ ’coverage’ was then used to quantify the presence of these genes in each sample and to calculate the percentage of reads mapped to these genes per sample.

### Feature Types

As described above, we used the 16S and shotgun metagenomic reads from our samples to generate five datasets, each consisting of a single feature type: 16S rRNA gene amplicon ASVs, species, strains, MAGs, and genes. The 16S ASVs are the only feature type that was generated using amplicon-based methods. The rest were based on the shotgun metagenomic reads. Within these remaining four, species and strains were read-based methods because they did not require assembly, while MAGs and genes were assembly-based, since they used contigs of assembled reads. We analyzed all five feature types to determine which method allowed us to minimize the spurious appearance of engraftment, while still having the sensitivity to detect true engraftment that may be happening.

### Assessing Engraftment

In order for us to call a feature “apparently engrafted”, it needed to meet three criteria: 1. It must be observed in the Donor B samples. 2. It must be absent from the patient’s pre-treatment sample. 3. It must be present in the patient’s post-treatment sample.

For all feature types, criterion 1 was met if the feature was observed at least once in any Donor B sample. The definitions of criteria 2 and 3 (absence and presence) depended on feature type. Our aim was to be quite strict with these criteria in order to minimize spurious identification of engraftment.

For the 16S ASVs, and strain and species marker feature types, a feature was deemed to be absent at baseline (criterion 2) if the patient’s pre-treatment sample contained that feature exactly 0 times. Even a single read assigned to a given feature in a baseline sample would remove it from consideration for engraftment in that patient. Criterion 3 (presence after treatment) was met if the feature’s relative abundance in the post-treatment sample surpassed a given threshold. The threshold varied between the three feature types, and in each case was chosen such that, when it was applied to Donor B’s own samples, it eliminated the bottom 20% of features from the merged datasets. The cutoffs were 1.3e04 relative abundance for 16S ASVs and strain markers, and 0.0072 for species markers. **Supp Figure 1** shows the presence vs. relative abundance histograms used to determine these cutoffs.

We additionally used the StrainPhlAn strain-sharing workflow^25^ to assess engraftment of strains. In this case, a strain was considered absent from a sample if its species-level genomic bin (SGB) pairwise distance was not below the pre-computed similarity threshold49 with any Donor B sample, and was considered present if its SGB pairwise distance was below the threshold. The results of this analysis can be found in the supplemental material.

For assembly-based feature types (genes and MAGs), presence and absence were determined by the proportion of the sequence with at least 1x coverage, rather than abundance. In order to identify reasonable cutoffs for MAGs, we mapped Donor B’s reads back to the database of Donor B assemblies, and then created a histogram showing the frequencies with which MAGs were covered at least 1x over different proportions of their length. We observed a u-shape in this histogram, where approximately 50% of the MAGs in each sample were covered 1x over at least 75% of their length, most intermediate values of coverage were present at very low frequency, and there was another peak of varying height when there was at least 1x coverage over less than 20% of the MAG’s length. We reason that MAGs that were present in a given sample are represented by the peak above 75% coverage, while MAGs that are absent from that sample can be seen in the lower peak. This threshold aligns with the theoretical minimum detectable genome overlap proposed by Lander and Waterman (1988)^58^. We called a MAG absent in a baseline sample if it was covered at least 1x over less than 20% of its length, and present in a post-treatment sample if it was covered at least 1x over at least 75% of its length. For genes, we used a 25% coverage criterion to define a gene as absent, and required its 1x coverage to be over at least 90% of its length in order to call it present after treatment. Both cutoffs were raised relative to the MAG cutoffs to allow for the fact that different genes which happen to share similar motifs can have quite similar DNA sequences over a substantial proportion of their lengths. Because we did not have empirical support for the gene cutoffs, we conducted a sensitivity analysis to ensure that our results were robust to changes in the absence and presence cutoffs (see supplemental results).

Within each patient, any features that failed to meet even one of the three criteria were removed from consideration for engraftment.

### Statistical Analysis of Engraftment

Once apparently engrafted features were identified in all samples across all feature types, we sought to test whether patients who received FMT had higher levels of apparent engraftment than those who received placebo treatment. We constructed three test statistics, all of which we tested using the same permutation test. These three test statistics were the differences between FMT and placebo groups in: 1. the number of features engrafted at least once in the given treatment group (T_1_), 2. the number of engraftment events in that occurred in each treatment group (i.e. the count of engrafted features, weighted by the number of patients each feature was engrafted in) (T_2_), and 3. the number of engraftment events weighted by the number of patients each event occurred in (.e. the count of engrafted features, weighted by the square of the number of patients each feature was engrafted in) (T_3_). The reason for this third test statistic was that we reasoned that a feature that appears engrafted in multiple patients is less likely to have been a spurious signal of engraftment, and so we wanted to up-weight those features based on the number of individuals they were engrafted in. Looking at the graphs in **Figure 2 (A1, B1, C1), 4(A1), and 5(A1**), the test statistics can be calculated as follows, where x is the number of patients a feature is engrafted in, and f(x) is the number of features engrafted in exactly that many patients:

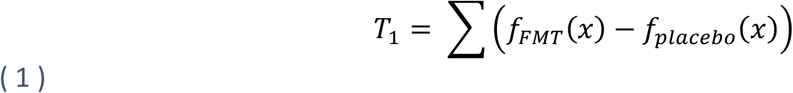

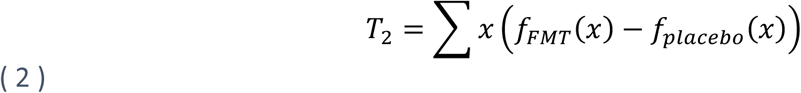

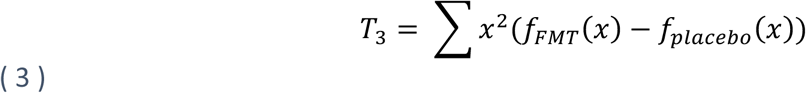

Permutation tests were conducted by permuting the group membership of each patient 1,999 times, and for each permutation calculating the number of engrafted features, engraftment events, and weighted engraftment events in each group, and taking their difference. The 2,000^th^ value in each permuted null distribution was the observed value. Two-tailed p-values were calculated by finding the quantile of the observed test statistic in the null distribution, taking its distance from 0 or 1, whichever was smaller, and doubling that value. This same method was used to compare the amounts of apparent engraftment in responder-vs. non-responder-patients within the FMT treatment group **(Figure 2(A3, B3, C3), 4(A3), and 5(A3)).**

Once count tables or coverage values had been calculated for a given feature type, all data organization and analysis code were written in R v4.2.0 using the tidyverse^59^ collection of packages. Figures were generated in R using ggplot2 and tidytext and refined in Inkscape. R scripts are available at https://github.com/SShekarriz/UCFMT1.

### Data and code availability

Shotgun metagenomic sequences used in this study are available via the SRA under BioProject number PRJNA1220173. These samples include fecal samples from donor B and patients with ulcerative colitis, as well as plate pool metagenomes from donor B. All code used for figure generation and statistical analyses is available at https://github.com/SShekarriz/UCFMT1.

## Acknowledgements

This work was supported by grants from Crohn’s Colitis Canada, and the Inflammation, Microbiome, and Alimentation: Gastrointestinal and Neuropsychiatric Effects (IMAGINE) Network CIHR grant. M.G.S. is supported by a Canada Research Chair. During some of this work S.S. was supported by Mitacs Elevate Fellowship and F.J. Whelan was supported by an Anne McLaren Fellow funded by the University of Nottingham and a Marie Skłodowska-Curie Individual Fellowship (grant agreement number 793818). The authors would like to thank Dr. Ben Bolker and the McMaster biodata lunch group, as well as Dr. Kevin Purbhoo, for their valuable insight and suggestions in the development of our statistical methods.

## Supplementary Results

### Culture-enriched metagenomics improves the quality of de novo assembly and taxonomic binning

It is often challenging to determine if the donor is responsible for the observed microbial changes after FMT or if they instead represent population restructuring within the patient’s own microbiota. Assembly-based metagenomic approaches provide better resolution, and may be more suitable than amplicon- or read-based metagenomic techniques for tracking strain- or gene-level changes. However, the quality of genome assembly and taxonomic binning of metagenomic assemblies can be poor^60^ and recovering low-abundance taxa is challenging^37^. Previous studies have shown that the human microbiota is culturable^26,28,29,62^, and that a combination of direct (culture-independent) sequencing and sequencing after comprehensive culturing can result in increased observed microbial diversity^26,29,63^.

We hypothesized that co-assembling sequences from direct and culture-enriched metagenomic sequencing (CEMG) of the Donor B microbiota would improve gene and metagenome assembled genome (MAG) recovery from this donor compared to the commonly used direct metagenomic (DMG) approach. To test this hypothesis, we conducted shotgun metagenomic sequencing on a previously cultured^26^ fecal sample from Donor B (**Figure 1**). We then compared the assemblies and microbial binning qualities achieved from DMG vs CEMG sequencing for this single fecal sample.

DMG resulted in 35,577 assembled contigs (> 2.5 Kb) accounting for ∼340 Mb of the assembled sequences, whereas the same number of contigs in CEMG captured ∼620 Mb. In addition to providing more contigs, the CEMG method produced longer contigs, specifically (**Supp Figure 2A**). These longer contigs enhanced gene predictions and generated more (132 vs. 49) metagenome-assembled genomes (MAGs; > 70% complete and < 10% contamination; see Methods). To assess the specificity of the two approaches, we mapped raw reads from both DMG and CEMG to the assembled MAGs from both assemblies. CEMG provided 83 more MAGs than DMG; however, most of these additional MAGs were well covered by the DMG reads, indicating that their presence is not due to contamination, but instead increased representation in the sequencing (**Supp Figure 2B**). The additional MAGs from CEMG sequencing are not derived from any particular taxonomic groups, but instead represent multiple families in proportion to their abundance in the DMG approach (**Supp Figure 2B, C**).

To examine the quality of assembled MAGs, we selected 40 homologous MAGs from the DMG and CEMG assemblies based on their position in a maximum-likelihood phylogenetic tree of core genome alignment (data not shown), and compared their genome sizes. We found that 24 out of the 40 MAGs (60%) were longer in the CEMG assemblies than in DMG (**Supp Figure 2D**). We concluded that CEMG sequencing enhances the assembly of genes and MAGs for intestinal microbiota and set to use this approach to investigate microbial engraftment following FMT.

### Gene engraftment sensitivity analysis

We conducted a sensitivity analysis of our presence and absence cutoffs for the gene data set, to ensure that the pattern we observed was not a result of the specific values we chose for absence (<=20% covered) or presence (>=90% covered). We tested all combinations of absence cutoffs ranging from 0% to 40% and presence cutoffs ranging from 70% to 100%, with step sizes of 5 percentage points. The results of the analysis were remarkably consistent across all combinations of values (Supp Figure 4 Supp Figure 5). We also investigated the distribution of coverage values in our patient samples (shown in Supp Figure 6). The bimodality of this distribution demonstrates that there is not much ambiguity in these data. Only a small proportion of genes are partially covered, compared to the number of genes that are covered at ∼0% or ∼100%, further supporting the robustness of our analysis to the specific presence and absence cutoffs.

### Strain-sharing inferences using StrainPhlAn4

508 SGBs detected in DonorB samples (n = 4) with relative abundance ≥ 0.001% were used to generate phylogenetic trees using StrainPhlAn4. We were able to construct SNV profiling for 169 out of 508 species-level markers (SGBs) with options for increased sensitivity (see Methods). Using pre-computed species-specific strain identity thresholds from Valles-Colomer et al. (2023)^25^, we identified 54 unique SGBs that showed engraftment in at least one patient at the strain level via the strain-sharing inference pipeline. Of those, 52 were engrafted only FMT samples, one was engrafted only a placebo sample, and one was engrafted in samples from both groups (**Supp Figure 7**). Counting how many SGBs were engrafted in multiple patients was a challenge. In **Supp Figure 7**, we counted an SGB as multiply engrafted if the pairwise distance between any donor sample and any patient sample was below the threshold for more than one patient sample. However, this does not guarantee that the specific strain shared between the donor and one patient is the same as the strain shared between the donor and another patient. To investigate this, we checked the between-patient distances against the pre-computed thresholds for the multiply engrafted SGBs. We found that about half the time, the between-patient distances were below the threshold, indicating that the patients were sharing a strain (data not shown). This categorization is further complicated by the continuous nature of the data, which means that strain-sharing may not be a transitive property of samples. For example (**Supp Figure 7**, inset), we found one SGB that was shared between donor samples and three patient samples, pt25, pt10, and pt62. Although the pairwise distances between pt25 and the other two samples were below the threshold, the pairwise distance between pt10 and pt62 was not.

We identified 252 unique SGBs engrafted in >=1 patients at the species level compared to 54 via strain transmission pipeline. These results suggest that the strain-sharing inference method considers only a small subset of SGBs for strain-sharing inference because it requires higher abundance of these SGBs to generate phylogenetic alignments across samples and markers (**Supp Figure 8).**

### Engrafted taxonomy

We investigated whether there is a consistent pattern of engrafted taxa across feature types. Although there is some noise, broadly speaking taxa that are engrafted at a high rate in one feature type are engrafted at a high rate in other feature types, and those that are not engrafted or minimally engrafted in one feature type are also minimally engrafted in other feature types. Supp Figure *9* shows this pattern.

## Supplementary Figures

## Supplementary Tables

**Supp Figure 1.**
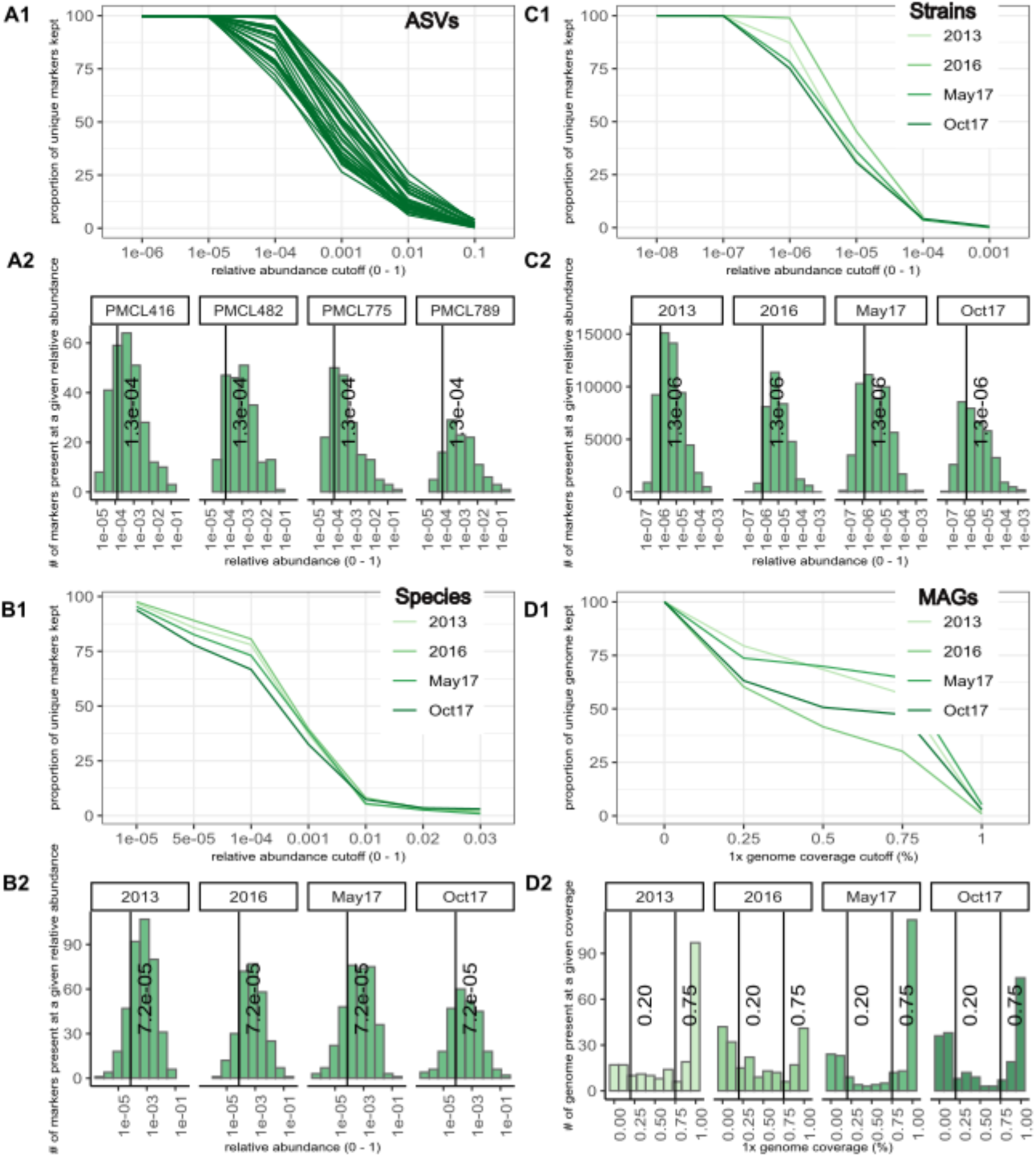
Profiles of features in Donor B samples, for determining engraftment cutoffs. Panels (A – D)1 show the proportion of unique features retained (y-axis) after all features below a given cutoff (x-axis) are removed. In panels A (16S ASVs), B (species markers), and C (strain markers), the cutoffs are relative abundances. In panel D (MAGs), the cutoffs are the proportion of the genome covered by at least one read. Panels (A – D)2 are histograms of the frequencies of various relative abundances (A – C) or 1x coverages (D). Vertical black lines in A2, B2, and C2 show our chosen cutoffs to define a feature as present, in both cases removing the bottom 20% of the density of the histogram. Vertical black lines in D2 show our chosen cutoffs for both absence and presence, removing the distributions’ two peaks as described in the Methods section.

**Supp Figure 2.**
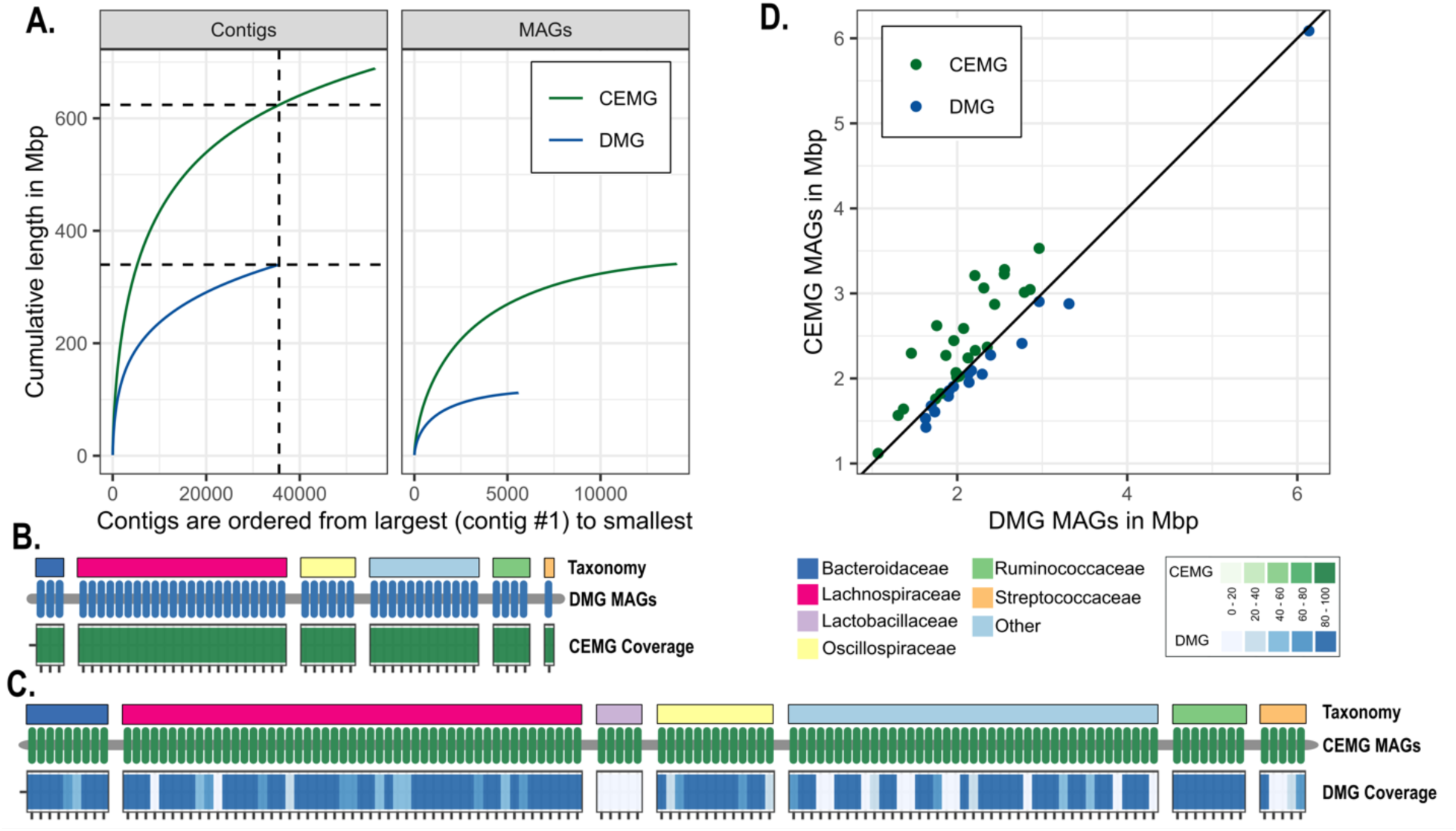
Comparison of culture enriched (CEMG) and direct metagenomics (DMG) for Donor B. **A.** The cumulative lengths of assembled contigs >2.5 kb for all contigs, and for contigs belonging to MAGs. Vertical dashed line represents the number of contigs in the DMG assembly, horizontal lines represent the total assembly size at that number of contigs for DMG and CEMG respectively. B. DMG and CEMG read mapping coverage (mean MAG coverage at least 1X) of the MAGs (n=49) assembled via DMG. The top colour bar shows the family-level taxonomy of the MAGs. C. CEMG and DMG read mapping coverage for the MAGs (n=132) assembled via CEMG. D. Genome sizes among homologous MAGs in CEMG and DMG.

**Supp Figure 3.**
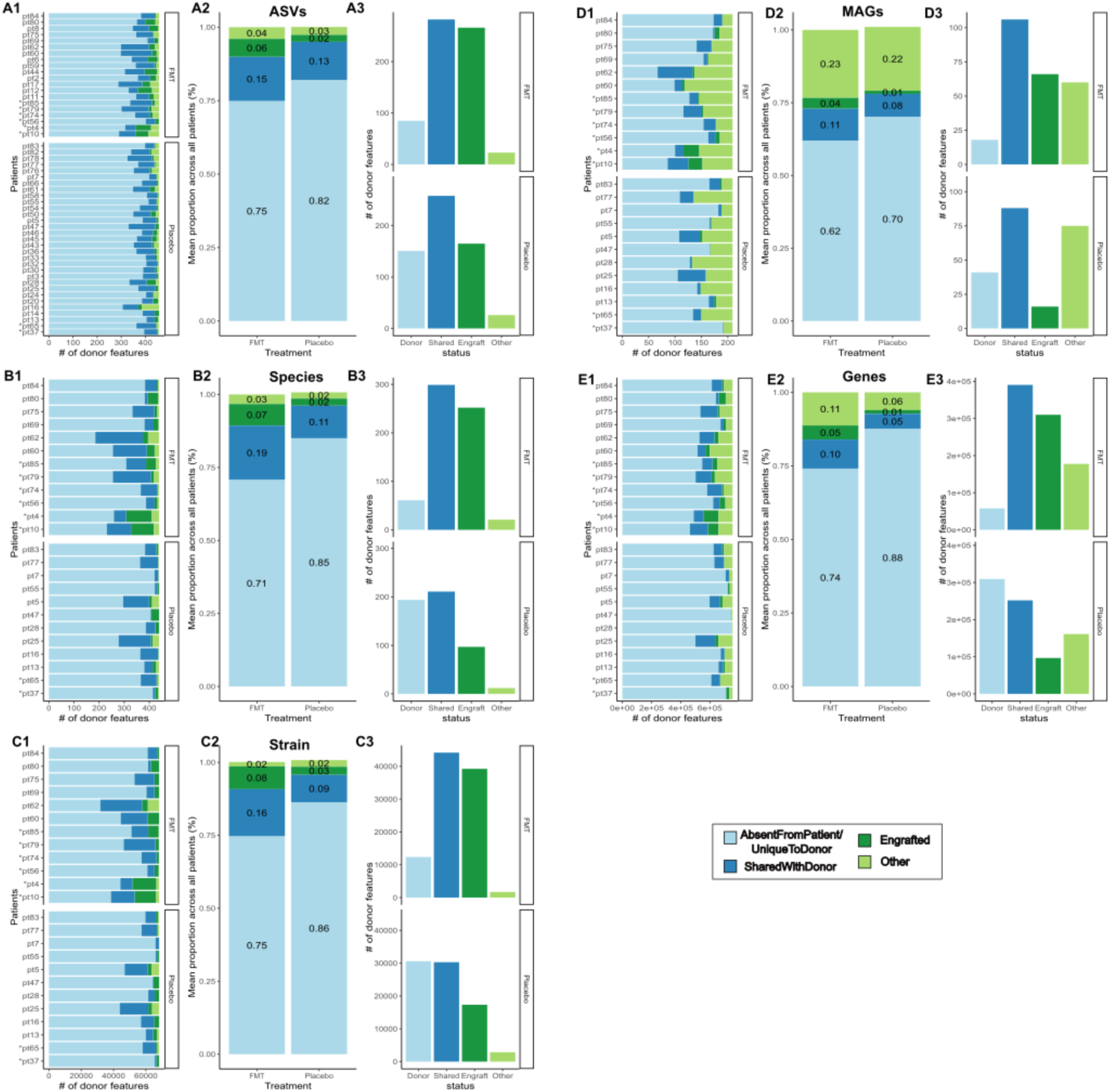
Proportion of Donor B’s features that are engrafted, shared between donor and recipient, or only found in donor. Panels A(1 – 3) concern the ASV feature type, panels B(1 – 3) concern the strain feature type, panels C(1 – 3) concern the species feature type, panels D(1 – 3) concern the MAG feature type, and panels E(1 – 3) concern the gene feature type. Panels (A – E)1: For each patient, we calculated the proportion of Donor B features of a given feature type that were absent (as defined by the absence cutoff of our engraftment criteria) from both the pre- and post-treatment samples (light blue: Donor), were already present (as defined by our presence cutoff ditto) in the pre-treatment sample (dark blue: Shared), met the criteria for engraftment (dark green: Engraft), or did not meet any of these criteria (light green: Other). In almost every case, the majority of Donor B’s features were absent from the patient, and the ‘Other’ category contained the fewest features. Panels (A – E)2: the mean values of the categories in (A – E)1 across all patients within a given feature type. Again showing that, within any individual patient, most Donor B features are absent from that patient at all times. Panels (A – E)3: Taking all patients in a given treatment arm (FMT or Placebo) together, features are counted in the light blue (Donor) category if they are absent from both timepoint samples from **all** patients. Features are counted in the Shared (dark blue) and Engraft (dark green) categories if they meet the corresponding above-define criteria in **any** patient (meaning some features may be counted in both categories). Features that do not meet any of the above criteria are included in the Other (light green) bar. In all cases, the Donor category contains more features in the Placebo group than the FMT group; however, the gene feature type shows the largest proportional difference. In all cases, the FMT group includes more Engraft features than the Placebo group. The MAG and gene features types show the largest proportional difference.

**Supp Figure 4.**
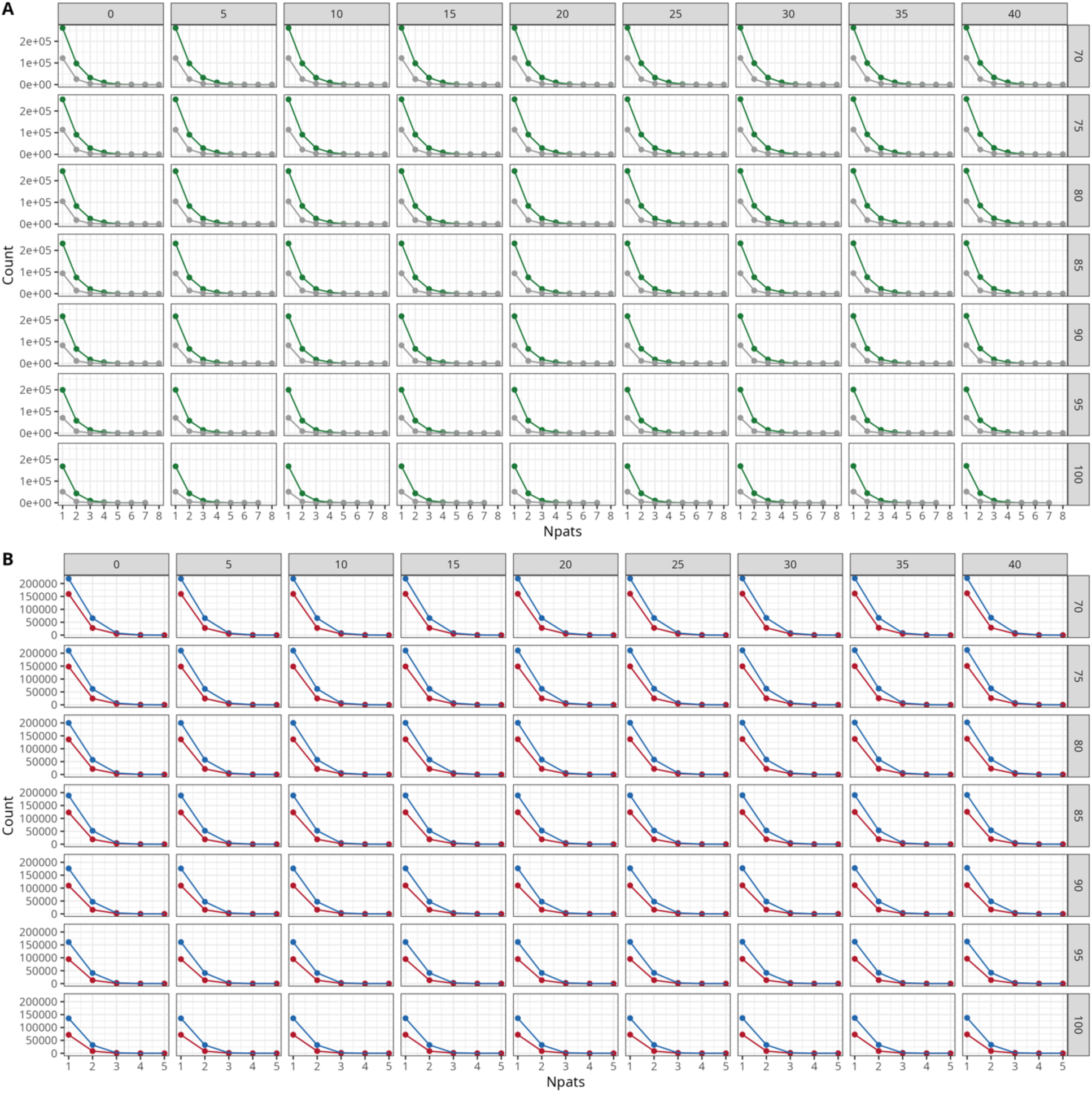
Number of apparent engraftment events in increasing numbers of patients. Columns are absence cutoffs ranging from 0% - 40%, rows are presence cutoffs ranging from 70% - 100%. A. apparent engraftment events in all FMT patients (green) and Placebo patients (grey). B. apparent engraftment events in FMT Remission patients (blue) and FMT No Remission patients (red).

**Supp Figure 5.**
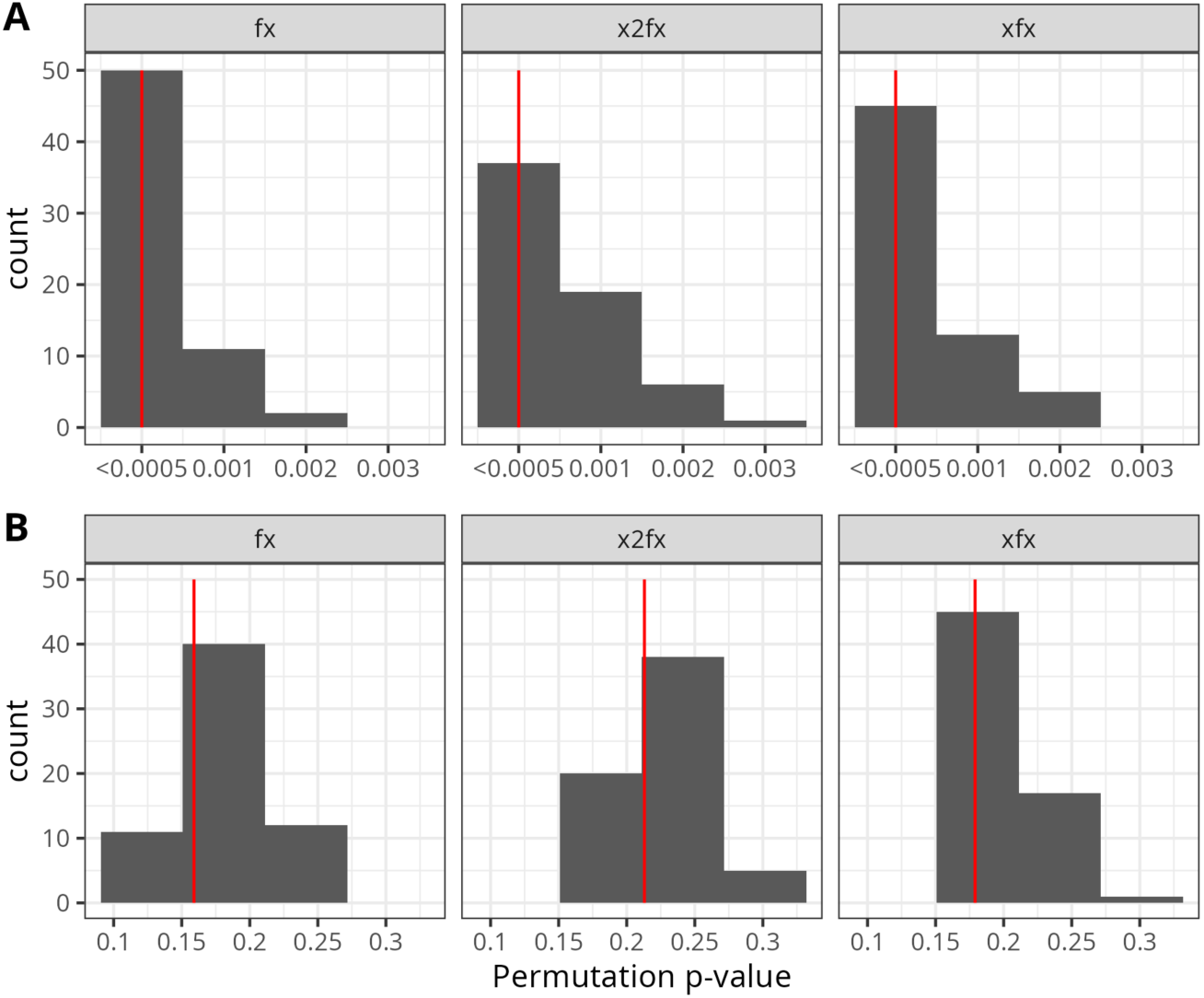
Histograms of the permutation test p-values calculated across all absence and presence cutoff values. The red line shows the p-value for the absence and presence cutoffs used in our analysis. This p-value is at or near the mode of the distribution for all statistical tests. A. p-values for FMT vs. Placebo samples. B. p-values for FMT Remission vs. FMT No Remission samples.

**Supp Figure 6.**
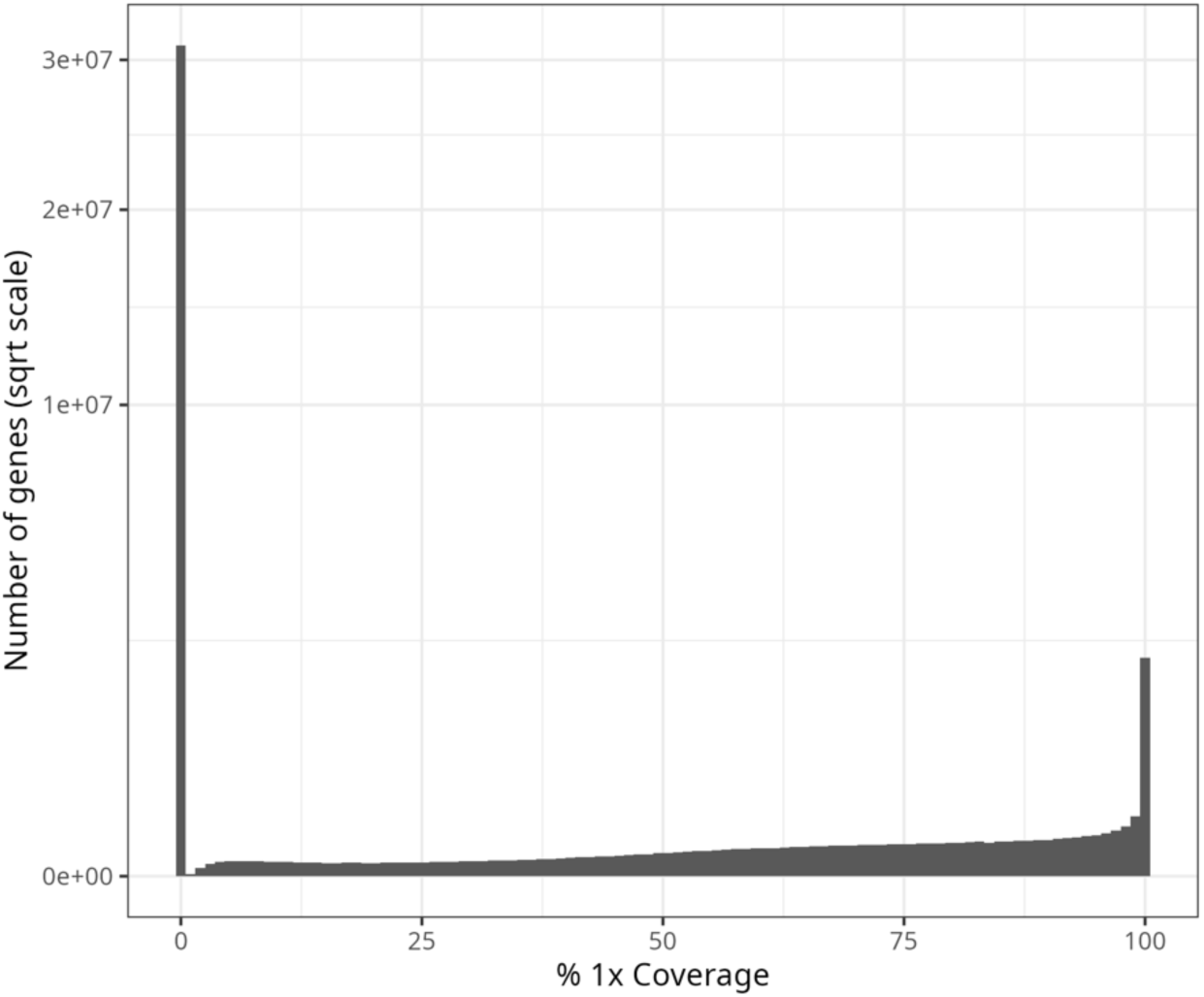
Histogram of the number of genes at given levels of 1x coverage across their lengths. The distribution is strongly bimodal, with the majority of its density near 0 or 100%. The y-axis is shown on the square-root scale so that intermediate coverage values are visible.

**Supp Figure 7.**
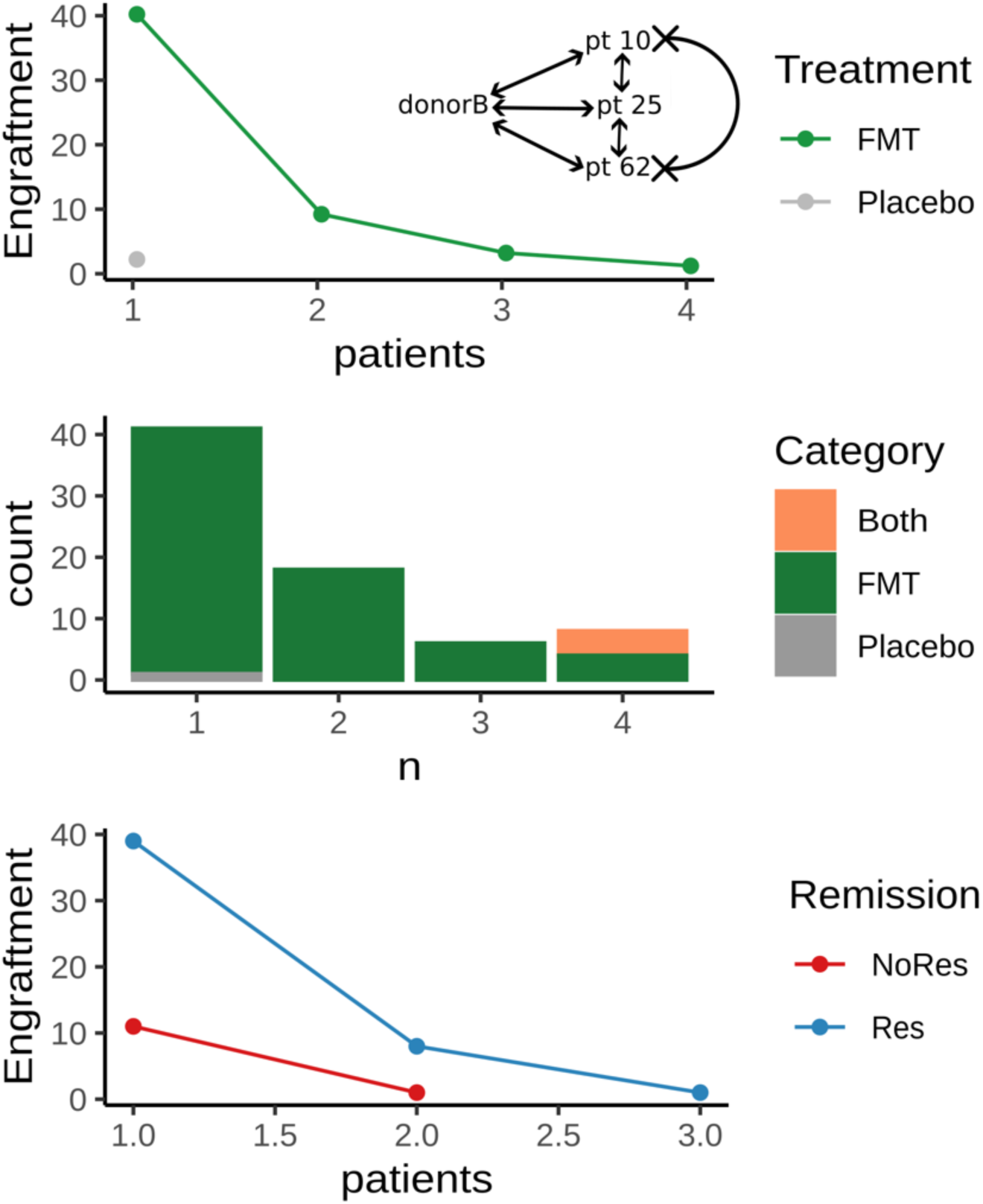
Comparison of rates of apparent engraftment between groups. Detection of apparently engrafted strains according to the StrainPhlAn strain-sharing method. Top panel: Comparison of the number of apparently engrafted features observed in FMT vs. placebo samples. The x-axis shows the number of patients a feature is apparently engrafted in, and the y-axis shows the number of features observed to be apparently engrafted in exactly that many patients within FMT (green) or placebo (grey) samples. Middle panel: Counting the number of apparently engrafted features that are unique to FMT samples (green), placebo samples (grey), or present in both (orange). The x-axis shows the number of patients a feature is apparently engrafted in, and the y-axis shows the number of features observed apparently engrafted in exactly that many patients. Bottom panel: The number of apparently engrafted features in FMT samples that are present in patients who responded to FMT treatment (blue lines) and did not respond (red lines). The inset diagram in the top panel demonstrates the problem of non-transitivity of strain sharing, where two samples can meet the strain-sharing threshold with the same third sample, but not with each other.

**Supp Figure 8.**
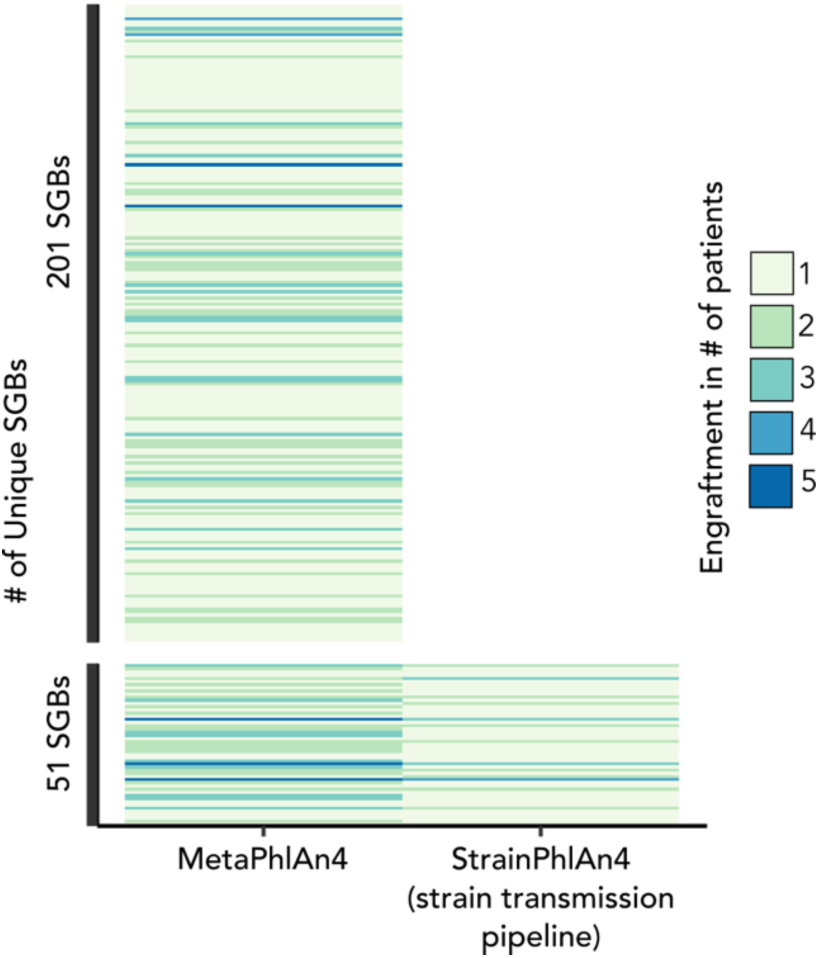
Comparison of the engrafted SGBs in increasing number of patients at species-level (MetaPhAn4) and stain marker-level using strain transmission pipeline (StrainPhlAn4).

**Supp Figure 9.**
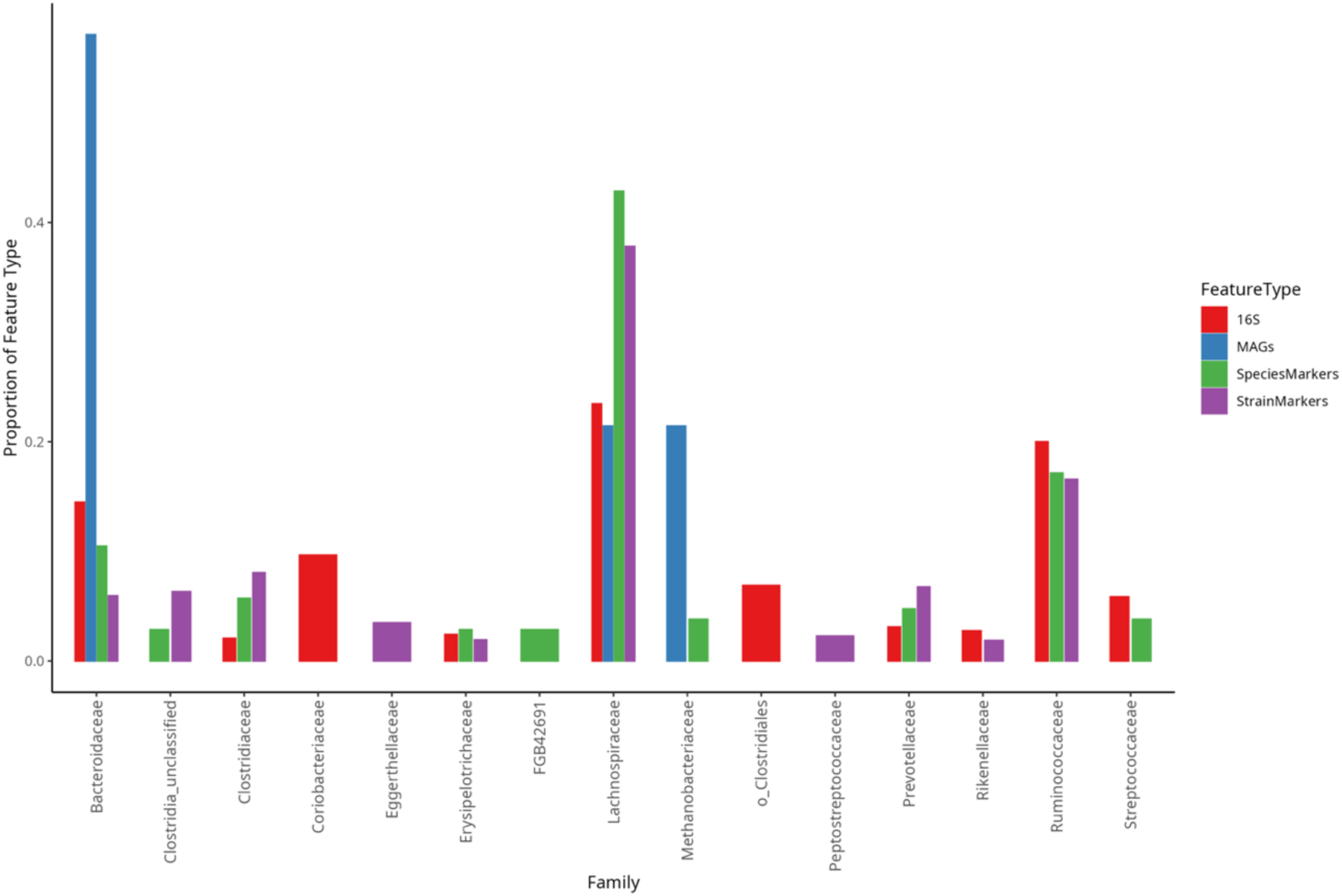
Comparison of Engrafted Microbial Families Across Feature Types. The proportion of engraftment events observed in FMT samples for each feature type, categorized by microbial family. Up to the 10 most common families per feature type are included, with features lacking taxonomic assignments excluded. The notably tall bars for MAGs likely reflect the limited number of detected engrafted MAGs.

**Supplementary Table 1:**
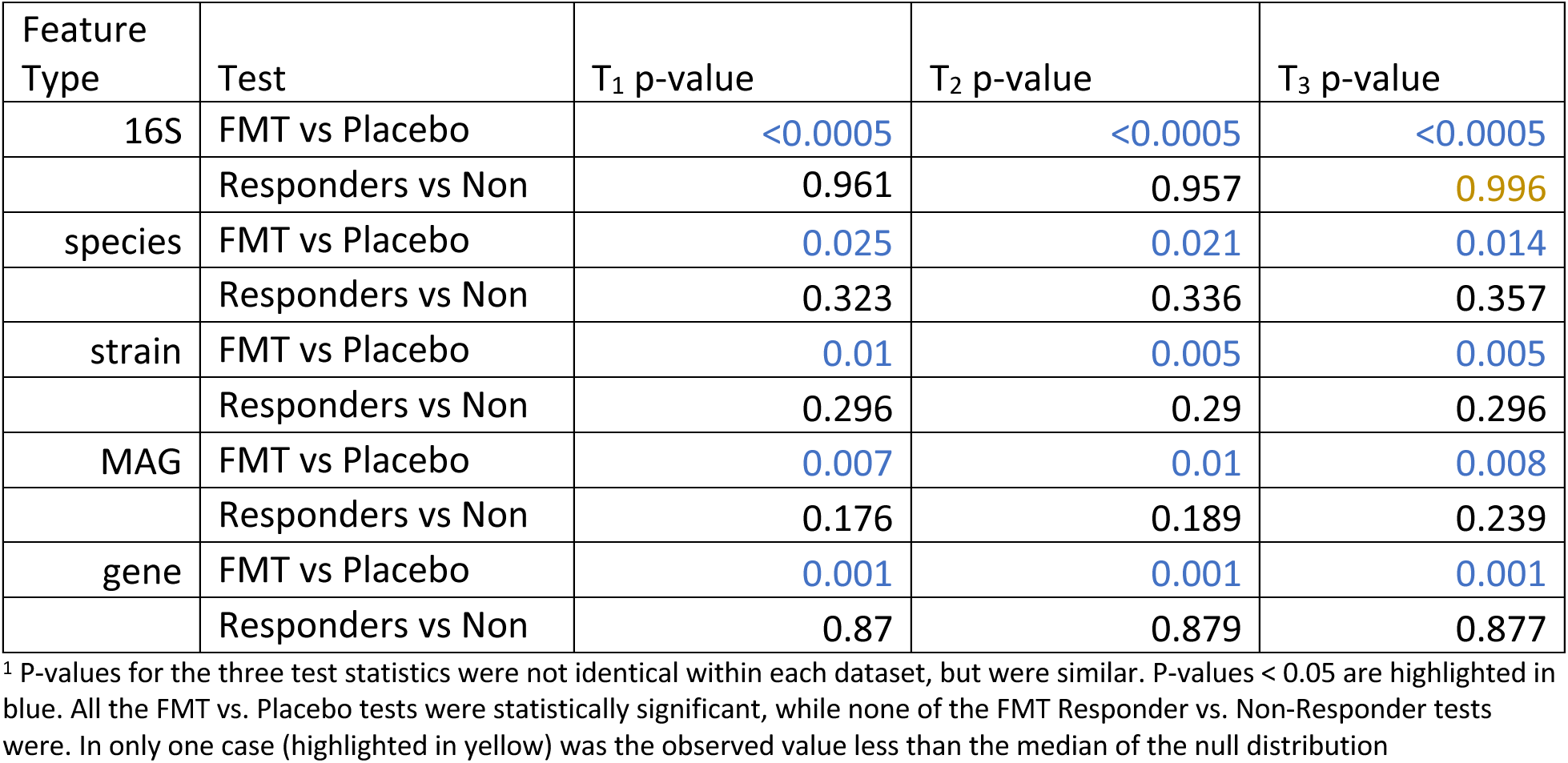
Permutation-test p-values for all three test statistics.^1^.

**Supplementary Table 2:** Number of features apparently engrafted, by number of patients that feature was engrafted in for each type of analysis.

| Feature Type | Condition | # Patients | Feature Count |
| --- | --- | --- | --- |
| 16S | Placebo | 1 | 81 |
|  |  | 2 | 35 |
|  |  | 3 | 29 |
|  |  | 4 | 11 |
|  |  | 5 | 4 |
|  |  | 6 | 3 |
|  |  | 7 | 1 |
|  |  | 8 | 0 |
|  |  | 9 | 0 |
|  |  | 10 | 1 |
|  | FMT (all) | 1 | 109 |
|  |  | 2 | 79 |
|  |  | 3 | 40 |
|  |  | 4 | 25 |
|  |  | 5 | 10 |
|  |  | 6 | 2 |
|  |  | 7 | 0 |
|  |  | 8 | 1 |
|  |  | 9 | 0 |
|  |  | 10 | 0 |
|  | FMT (responder) | 1 | 118 |
|  |  | 2 | 57 |
|  |  | 3 | 33 |
|  |  | 4 | 9 |
|  |  | 5 | 3 |
|  |  | 6 | 0 |
|  |  | 7 | 1 |
|  | FMT (non-responder) | 1 | 96 |
|  |  | 2 | 33 |
|  |  | 3 | 2 |
|  |  | 4 | 0 |
|  |  | 5 | 0 |
|  |  | 6 | 0 |
|  |  | 7 | 0 |
| species | Placebo | 1 | 80 |
|  |  | 2 | 14 |
|  |  | 3 | 3 |
|  |  | 4 | 0 |
|  |  | 5 | 0 |
|  | FMT (all) | 1 | 158 |
|  |  | 2 | 63 |
|  |  | 3 | 24 |
|  |  | 4 | 2 |
|  |  | 5 | 5 |
|  | FMT (responder) | 1 | 146 |
|  |  | 2 | 49 |
|  |  | 3 | 5 |
|  | FMT (non-responder) | 1 | 90 |
|  |  | 2 | 14 |
|  |  | 3 | 4 |
| strain | Placebo | 1 | 13,080 |
|  |  | 2 | 3,338 |
|  |  | 3 | 750 |
|  |  | 4 | 148 |
|  |  | 5 | 20 |
|  |  | 6 | 1 |
|  |  | 7 | 0 |
|  |  | 8 | 0 |
|  | FMT (all) | 1 | 23,093 |
|  |  | 2 | 10,316 |
|  |  | 3 | 4,008 |
|  |  | 4 | 1,229 |
|  |  | 5 | 433 |
|  |  | 6 | 129 |
|  |  | 7 | 29 |
|  |  | 8 | 1 |
|  | FMT (responder) | 1 | 21,322 |
|  |  | 2 | 7,281 |
|  |  | 3 | 1,438 |
|  |  | 4 | 162 |
|  | FMT (non-responder) | 1 | 14,737 |
|  |  | 2 | 3,084 |
|  |  | 3 | 620 |
|  |  | 4 | 51 |
| MAG | Placebo | 1 | 14 |
|  |  | 2 | 2 |
|  |  | 3 | 0 |
|  |  | 4 | 0 |
|  |  | 5 | 0 |
|  | FMT (all) | 1 | 45 |
|  |  | 2 | 17 |
|  |  | 3 | 3 |
|  |  | 4 | 0 |
|  |  | 5 | 1 |
|  | FMT (responder) | 1 | 39 |
|  |  | 2 | 15 |
|  |  | 3 | 1 |
|  | FMT (non-responder) | 1 | 18 |
|  |  | 2 | 0 |
|  |  | 3 | 1 |
| gene | Placebo | 1 | 83,282 |
|  |  | 2 | 11,775 |
|  |  | 3 | 1,472 |
|  |  | 4 | 142 |
|  |  | 5 | 9 |
|  |  | 6 | 0 |
|  |  | 7 | 0 |
|  |  | 8 | 0 |
|  | FMT (all) | 1 | 218,257 |
|  |  | 2 | 67,350 |
|  |  | 3 | 18,043 |
|  |  | 4 | 5,511 |
|  |  | 5 | 713 |
|  |  | 6 | 91 |
|  |  | 7 | 11 |
|  |  | 8 | 1 |
|  | FMT (responder) | 1 | 176,849 |
|  |  | 2 | 47,623 |
|  |  | 3 | 3,766 |
|  |  | 4 | 258 |
|  |  | 5 | 5 |
|  | FMT (non-responder) | 1 | 110,286 |
|  |  | 2 | 16,322 |
|  |  | 3 | 1,873 |
|  |  | 4 | 78 |
|  |  | 5 | 3 |

**Supplementary Table 3:** The number of features apparently engrafted uniquely in FMT or Placebo groups, or in both, by the total number of patients they appeared to be engrafted in.

| Feature Type | Group Engrafted | Total Patients | Count |
| --- | --- | --- | --- |
| 16S | FMT Only | 1 | 73 |
|  |  | 2 | 42 |
|  |  | 3 | 18 |
|  |  | 4 | 9 |
|  |  | 5 | 4 |
|  | Placebo Only | 1 | 31 |
|  |  | 2 | 8 |
|  |  | 3 | 4 |
|  |  | 4 | 2 |
|  | Both | 2 | 18 |
|  |  | 3 | 20 |
|  |  | 4 | 28 |
|  |  | 5 | 19 |
|  |  | 6 | 18 |
|  |  | 7 | 5 |
|  |  | 8 | 5 |
|  |  | 9 | 3 |
|  |  | 10 | 2 |
|  |  | 12 | 1 |
|  |  | 15 | 1 |
| species | FMT Only | 1 | 123 |
|  |  | 2 | 48 |
|  |  | 3 | 16 |
|  | Placebo Only | 1 | 25 |
|  |  | 2 | 6 |
|  |  | 3 | 1 |
|  | Both | 2 | 28 |
|  |  | 3 | 20 |
|  |  | 4 | 8 |
|  |  | 5 | 4 |
|  |  | 6 | 4 |
|  |  | 7 | 1 |
| strain | FMT Only | 1 | 16,872 |
|  |  | 2 | 6,594 |
|  |  | 3 | 2,343 |
|  |  | 4 | 552 |
|  |  | 5 | 138 |
|  |  | 6 | 67 |
|  |  | 7 | 20 |
|  |  | 8 | 1 |
|  | Placebo Only | 1 | 3,589 |
|  |  | 2 | 865 |
|  |  | 3 | 193 |
|  |  | 4 | 33 |
|  |  | 5 | 6 |
|  | Both | 2 | 4,618 |
|  |  | 3 | 4,053 |
|  |  | 4 | 2,245 |
|  |  | 5 | 1,078 |
|  |  | 6 | 474 |
|  |  | 7 | 154 |
|  |  | 8 | 26 |
|  |  | 9 | 3 |
| MAG | FMT Only | 1 | 39 |
|  |  | 2 | 16 |
|  |  | 3 | 3 |
|  | Placebo Only | 1 | 7 |
|  |  | 2 | 1 |
|  | Both | 2 | 5 |
|  |  | 3 | 2 |
|  |  | 6 | 1 |
| gene | FMT Only | 1 | 188,238 |
|  |  | 2 | 50,809 |
|  |  | 3 | 13,761 |
|  |  | 4 | 4,654 |
|  |  | 5 | 519 |
|  |  | 6 | 61 |
|  |  | 7 | 6 |
|  | Placebo Only | 1 | 39,487 |
|  |  | 2 | 4,672 |
|  |  | 3 | 555 |
|  |  | 4 | 35 |
|  |  | 5 | 2 |
|  | Both | 2 | 26,103 |
|  |  | 3 | 16,986 |
|  |  | 4 | 6,366 |
|  |  | 5 | 1,858 |
|  |  | 6 | 485 |
|  |  | 7 | 95 |
|  |  | 8 | 34 |
|  |  | 9 | 2 |

## References

1. Kappelman, M. D. et al. The prevalence and geographic distribution of Crohn’s disease and ulcerative colitis in the United States. Clin. Gastroenterol. Hepatol. Off. Clin. Pract. J. Am. Gastroenterol. Assoc. 5, 1424–1429 (2007).

2. de Souza, H. S. P. & Fiocchi, C. Immunopathogenesis of IBD: current state of the art. Nat. Rev. Gastroenterol. Hepatol. 13, 13–27 (2016).

3. Hindryckx, P., Jairath, V. & D’Haens, G. Acute severe ulcerative colitis: from pathophysiology to clinical management. Nat. Rev. Gastroenterol. Hepatol. 13, 654–664 (2016).

4. Talley, N. J. et al. An evidence-based systematic review on medical therapies for inflammatory bowel disease. Am. J. Gastroenterol. 106 **Suppl 1**, S2–25; quiz S26 (2011).

5. Moayyedi, P. et al. Fecal Microbiota Transplantation Induces Remission in Patients With Active Ulcerative Colitis in a Randomized Controlled Trial. Gastroenterology 149, 102–109.e6 (2015).

6. Peery, A. F. et al. AGA Clinical Practice Guideline on Fecal Microbiota-Based Therapies for Select Gastrointestinal Diseases. Gastroenterology 166, 409–434 (2024).

7. Khoruts, A., Staley, C. & Sadowsky, M. J. Faecal microbiota transplantation for Clostridioides difficile: mechanisms and pharmacology. Nat. Rev. Gastroenterol. Hepatol. 18, 67–80 (2021).

8. Khanna, S., Shin, A. & Kelly, C. P. Management of Clostridium difficile Infection in Inflammatory Bowel Disease: Expert Review from the Clinical Practice Updates Committee of the AGA Institute. Clin. Gastroenterol. Hepatol. Off. Clin. Pract. J. Am. Gastroenterol. Assoc. 15, 166–174 (2017).

9. Feng, J. et al. Efficacy and safety of fecal microbiota transplantation in the treatment of ulcerative colitis: a systematic review and meta-analysis. Sci. Rep. 13, 14494 (2023).

10. Pai, N. et al. Results of the First Pilot Randomized Controlled Trial of Fecal Microbiota Transplant In Pediatric Ulcerative Colitis: Lessons, Limitations, and Future Prospects. Gastroenterology 161, 388–393.e3 (2021).

11. Narula, N. et al. Systematic Review and Meta-analysis: Fecal Microbiota Transplantation for Treatment of Active Ulcerative Colitis. Inflamm. Bowel Dis. 23, 1702–1709 (2017).

12. Jaramillo, A. P. et al. Effectiveness of Fecal Microbiota Transplantation Treatment in Patients With Recurrent Clostridium difficile Infection, Ulcerative Colitis, and Crohn’s Disease: A Systematic Review. Cureus 15, e42120 (2023).

13. Liu, H., Li, J., Yuan, J., Huang, J. & Xu, Y. Fecal microbiota transplantation as a therapy for treating ulcerative colitis: an overview of systematic reviews. BMC Microbiol. 23, 371 (2023).

14. Smillie, C. S. et al. Strain Tracking Reveals the Determinants of Bacterial Engraftment in the Human Gut Following Fecal Microbiota Transplantation. Cell Host Microbe 23, 229–240.e5 (2018).

15. Staley, C. et al. Durable Long-Term Bacterial Engraftment following Encapsulated Fecal Microbiota Transplantation To Treat Clostridium difficile Infection. mBio 10, 10.1128/mbio.01586-19 (2019).

16. Aggarwala, V. et al. Precise quantification of bacterial strains after fecal microbiota transplantation delineates long-term engraftment and explains outcomes. Nat. Microbiol. 6, 1309–1318 (2021).

17. Deng, Z.-L. et al. Engraftment of essential functions through multiple fecal microbiota transplants in chronic antibiotic-resistant pouchitis—a case study using metatranscriptomics. Microbiome 11, 269 (2023).

18. Wilson, B. C. et al. Strain engraftment competition and functional augmentation in a multi-donor fecal microbiota transplantation trial for obesity. Microbiome 9, 107 (2021).

19. Chen-Liaw, A. et al. Gut microbiota strain richness is species specific and affects engraftment. Nature 637, 422–429 (2025).

20. Podlesny, D. et al. Identification of clinical and ecological determinants of strain engraftment after fecal microbiota transplantation using metagenomics. Cell Rep. Med. 3, 100711 (2022).

21. Ianiro, G. et al. Variability of strain engraftment and predictability of microbiome composition after fecal microbiota transplantation across different diseases. Nat. Med. 28, 1913–1923 (2022).

22. Schmidt, T. S. B. et al. Drivers and determinants of strain dynamics following fecal microbiota transplantation. Nat. Med. 28, 1902–1912 (2022).

23. Dsouza, M. et al. Colonization of the live biotherapeutic product VE303 and modulation of the microbiota and metabolites in healthy volunteers. Cell Host Microbe 30, 583–598.e8 (2022).

24. Menon, R. et al. Multi-omic profiling a defined bacterial consortium for treatment of recurrent Clostridioides difficile infection. Nat. Med. 31, 223–234 (2025).

25. Valles-Colomer, M. et al. The person-to-person transmission landscape of the gut and oral microbiomes. Nature 614, 125–135 (2023).

26. Lau, J. T. et al. Capturing the diversity of the human gut microbiota through culture-enriched molecular profiling. Genome Med. 8, 72 (2016).

27. Browne, H. P. et al. Culturing of ‘unculturable’ human microbiota reveals novel taxa and extensive sporulation. Nature 533, 543–546 (2016).

28. Sibley, C. D. et al. Culture Enriched Molecular Profiling of the Cystic Fibrosis Airway Microbiome. PLOS ONE 6, e22702 (2011).

29. Whelan, F. J. et al. Culture-enriched metagenomic sequencing enables in-depth profiling of the cystic fibrosis lung microbiota. Nat. Microbiol. 5, 379–390 (2020).

30. Myles, I. A. et al. A method for culturing Gram-negative skin microbiota. BMC Microbiol. 16, 60 (2016).

31. Hilt, E. E. et al. Urine Is Not Sterile: Use of Enhanced Urine Culture Techniques To Detect Resident Bacterial Flora in the Adult Female Bladder. J. Clin. Microbiol. 52, 871–876 (2014).

32. Callahan, B. J. et al. DADA2: High-resolution sample inference from Illumina amplicon data. Nat. Methods 13, 581–583 (2016).

33. Blanco-Míguez, A. et al. Extending and improving metagenomic taxonomic profiling with uncharacterized species using MetaPhlAn 4. Nat. Biotechnol. 41, 1633–1644 (2023).

34. Meziti, A. et al. The Reliability of Metagenome-Assembled Genomes (MAGs) in Representing Natural Populations: Insights from Comparing MAGs against Isolate Genomes Derived from the Same Fecal Sample. Appl. Environ. Microbiol. 87, e02593–20 (2021).

35. Franzosa, E. A. et al. Gut microbiome structure and metabolic activity in inflammatory bowel disease. Nat. Microbiol. 4, 293–305 (2019).

36. Montrose, J. A., Kurada, S. & Fischer, M. Current and future microbiome-based therapies in inflammatory bowel disease. Curr. Opin. Gastroenterol. 10.1097/MOG.0000000000001027 doi:10.1097/MOG.0000000000001027.

37. Costello, S. P. et al. Effect of Fecal Microbiota Transplantation on 8-Week Remission in Patients With Ulcerative Colitis: A Randomized Clinical Trial. JAMA 321, 156–164 (2019).

38. Whelan, F. J. et al. The loss of topography in the microbial communities of the upper respiratory tract in the elderly. Ann. Am. Thorac. Soc. 11, 513–521 (2014).

39. Bartram, A. K., Lynch, M. D. J., Stearns, J. C., Moreno-Hagelsieb, G. & Neufeld, J. D. Generation of multimillion-sequence 16S rRNA gene libraries from complex microbial communities by assembling paired-end illumina reads. Appl. Environ. Microbiol. 77, 3846– 3852 (2011).

40. Martin, M. Cutadapt removes adapter sequences from high-throughput sequencing reads. EMBnet.journal 17, 10–12 (2011).

41. Quast, C. et al. The SILVA ribosomal RNA gene database project: improved data processing and web-based tools. Nucleic Acids Res. 41, D590–D596 (2013).

42. McMurdie, P. J. & Holmes, S. phyloseq: An R Package for Reproducible Interactive Analysis and Graphics of Microbiome Census Data. PLOS ONE 8, e61217 (2013).

43. Bolger, A. M., Lohse, M. & Usadel, B. Trimmomatic: a flexible trimmer for Illumina sequence data. Bioinformatics 30, 2114–2120 (2014).

44. Schmieder, R. & Edwards, R. Fast Identification and Removal of Sequence Contamination from Genomic and Metagenomic Datasets. PLOS ONE 6, e17288 (2011).

45. Nurk, S., Meleshko, D., Korobeynikov, A. & Pevzner, P. A. metaSPAdes: a new versatile metagenomic assembler. Genome Res. 27, 824–834 (2017).

46. Li, H. & Durbin, R. Fast and accurate short read alignment with Burrows–Wheeler transform. Bioinformatics 25, 1754–1760 (2009).

47. Kang, D. D. et al. MetaBAT 2: an adaptive binning algorithm for robust and efficient genome reconstruction from metagenome assemblies. PeerJ 7, e7359 (2019).

48. Parks, D. H., Imelfort, M., Skennerton, C. T., Hugenholtz, P. & Tyson, G. W. CheckM: assessing the quality of microbial genomes recovered from isolates, single cells, and metagenomes. Genome Res. 25, 1043–1055 (2015).

49. Eren, A. M. et al. Community-led, integrated, reproducible multi-omics with anvi’o. Nat. Microbiol. 6, 3–6 (2021).

50. Chaumeil, P.-A., Mussig, A. J., Hugenholtz, P. & Parks, D. H. GTDB-Tk: a toolkit to classify genomes with the Genome Taxonomy Database. Bioinformatics 36, 1925–1927 (2020).

51. Price, M. N., Dehal, P. S. & Arkin, A. P. FastTree 2 – Approximately Maximum-Likelihood Trees for Large Alignments. PLOS ONE 5, e9490 (2010).

52. Schwengers, O. et al. Bakta: rapid and standardized annotation of bacterial genomes via alignment-free sequence identification. *Microb*. Genomics 7, 000685 (2021).

53. Shen, W., Le, S., Li, Y. & Hu, F. SeqKit: A Cross-Platform and Ultrafast Toolkit for FASTA/Q File Manipulation. PLOS ONE 11, e0163962 (2016).

54. Li, H. et al. The Sequence Alignment/Map format and SAMtools. Bioinformatics 25, 2078–2079 (2009).

55. Camacho, C. et al. BLAST+: architecture and applications. BMC Bioinformatics 10, 421 (2009).

56. Steinegger, M. & Söding, J. MMseqs2 enables sensitive protein sequence searching for the analysis of massive data sets. Nat. Biotechnol. 35, 1026–1028 (2017).

57. Cantalapiedra, C. P., Hernández-Plaza, A., Letunic, I., Bork, P. & Huerta-Cepas, J. eggNOG-mapper v2: Functional Annotation, Orthology Assignments, and Domain Prediction at the Metagenomic Scale. Mol. Biol. Evol. 38, 5825–5829 (2021).

58. Lander, E. S. & Waterman, M. S. Genomic mapping by fingerprinting random clones: a mathematical analysis. Genomics 2, 231–239 (1988).

59. Wickham, H. et al. Welcome to the Tidyverse. J. Open Source Softw. 4, 1686 (2019).

60. Sieber, C. M. K. et al. Recovery of genomes from metagenomes via a dereplication, aggregation and scoring strategy. Nat. Microbiol. 3, 836–843 (2018).

61. Sczyrba, A. et al. Critical Assessment of Metagenome Interpretation-a benchmark of metagenomics software. Nat. Methods 14, 1063–1071 (2017).

62. Lewis, W. H., Tahon, G., Geesink, P., Sousa, D. Z. & Ettema, T. J. G. Innovations to culturing the uncultured microbial majority. Nat. Rev. Microbiol. 19, 225–240 (2021).

63. Forster, S. C. et al. A human gut bacterial genome and culture collection for improved metagenomic analyses. Nat. Biotechnol. 37, 186–192 (2019).

